# Gain of function in *linc-133* compensates for *daf-18/PTEN* loss and rescues survival

**DOI:** 10.1101/2023.11.20.567969

**Authors:** Wei Huang, Fangzhou Dai, Lu Zhang, Xue Yin, Zhi Qu, Shanqing Zheng

## Abstract

DAF-18, homology of PTEN, possess tumor suppressor activity. Loss of *daf-18* causes cell growth in L1 diapause *C. elegans* is well studied; however, the reason why survival is dramatically shortened is not well elaborated. We found that *linc-133* gain of function can fully restore the shortened survival caused by *daf-18* loss. When lipid phosphatase activity of DAF-18 is defective, the *linc-133* gain of function interacts with 14-3-3 proteins to obstruct DAF-16 translocation from the nucleus to the cytoplasm. However, the dysfunction of protein phosphatase activity of DAF-18 caused high levels of aggregated proteins. The *linc-133* gain of function can induce HSP chaperones to select and process the aggregated proteins for degradation through ubiquitination. Our work demonstrates that protein homeostasis controlled by the protein phosphatase activity of DAF-18 is the main factor affecting survival and identifies a new function of the *linc-133* gene, which can compensate for the loss of *daf-18*.

## Introduction

L1 arrest larvae of *Caenorhabditis elegans* (*C. elegans*) provides an ideal model in which to study the coordination of biological phenomena and gene regulation ^1–5^. PTEN has been identified as a tumor suppressor in humans ^6^. The *C. elegans* PTEN homolog DAF-18 also possesses tumor suppressor-like properties. Wild-type N2 L1-arrested worms can live for approximately 20 days in sterilized liquid medium with no cell division. Loss of *daf-18* can break this diapause and shorten survival drastically ^2,3,7,8^. A variety of genes were reported to be related to L1 survival and the postembryonic cell developmental events that occur in *daf-18* mutants ^2,4,5,9,10^; however, we found that none of them could fully rescue the survival of *daf-18* worms during L1 arrest. DAF-18/PTEN is a dual-specificity phosphatase that has lipid and protein phosphatase activities ^11–13^. Its better-known lipid phosphatase activity, which controls the insulin/insulin-like pathway (IIS), was reported to play a pivotal role in regulating cell development during L1 arrest ^2,4,8,14^. However, genetic analyses show that the effect of *daf-18* loss on L1 arrest survival may not only occur through the regulation of PI3K-DAF-16/FOXO signaling in IIS ^2,5,7,8,10^. How downstream regulatory events control L1 arrest survival when *daf-18* is lost still needs to be explored.

In this work, we demonstrated that both the lipid and protein phosphatase activities of DAF-18 are needed to fully rescue the shortened survival caused by *daf-18* loss in L1-arrested worms. The high internal aggregated protein level caused by the dysfunction of protein phosphatase activity of DAF-18 may be the main reason why the worms have a shorter survival. By using a traditional forward genetic method ^15^, we identified that a gain-of-function mutation in *linc-133* can extend the survival of worms during L1 arrest and fully recover the survival of *daf-18* worms to that of wild-type worms. This new *linc-133* mutation is capable of compensating for the consequences of the functional defects in DAF-18 lipid and protein phosphatase activities.

## Results

### Both lipid and protein phosphatase activities of DAF-18/PTEN are needed to maintain survival

DAF-18/PTEN is well known for its lipid phosphatase activity, which plays an opposite role to AGE-1/PI3K in dephosphorylating phosphatidylinositol 3,4,5-trisphosphate (PIP3) to PIP2 ^16–18^ to negatively regulate the function of DAF-16/FOXO in IIS ^19^. We found that disrupting *age-1* can partially rescue *daf-18* survival during L1 arrest (Fig. 1a) and that the loss of *daf-16* also reduced the survival of L1-arrested worms relative to that of N2 worms (Fig. 1b); however, the survival of the *daf-16; daf-18* double mutant was even shorter than that of the *daf-16* single mutants (Fig. 1b), suggesting that *daf-18* also acts through mechanisms other than PI3K-AKT-DAF-16/FOXO to regulate L1 arrest survival. To confirm this speculation, we tested whether activated DAF-16 can fully rescue the shortened survival of *daf-18*. We found that DAF-16 activation, through inhibiting AKT, only partially rescued the survival of *daf-18* worms (Fig. 1c). Moreover, blocking IIS signaling by *daf-2* can rescue the short sruvival of *ins-3* (oe), an agonist gene of IIS, worms, but failed to rescue the short survival of *daf-18* worms (Fig. 1d). These results suggested that the DAF-16/FOXO is not the main target regulated by DAF-18 to control L1 arrest survival. DAF-18/PTEN also has protein phosphatase activity, which is independent of the PI3K-DAF-16/FOXO pathway ^12,20–22^. To identify which phosphatase activity of DAF-18 was required for supporting L1 arrest survival, three variants of DAF-18 that correspond to known human PTEN variants were constructed and tested for survival rescuing activity ^13^ (Fig. 1e). The results showed that the protein phosphatase-defective (DAF-18 with D137A) and lipid phosphatase-defective (DAF-18 with G174E) constructs both partially rescued the short survival of *daf-18* worms (Fig. 1f-g). DAF-18 (C169S) abolishes both lipid and protein phosphatase activity and is unable to rescue *daf-18* survival during L1 arrest (Fig. 1h). We also generated a *daf-18(D137A)* mutation strain by using CRISPR/Cas9. Interestingly, *daf-18(D137A)* worms lived only approximately 5 days during L1 arrest (Fig. 1i). These results confirm that survival of L1 arrest requires both the lipid and protein phosphatase activities of DAF-18 and suggest that DAF-18 protein phosphatase activity may play a pivotal role in supporting survival during L1 arrest.

**Figure 1.**
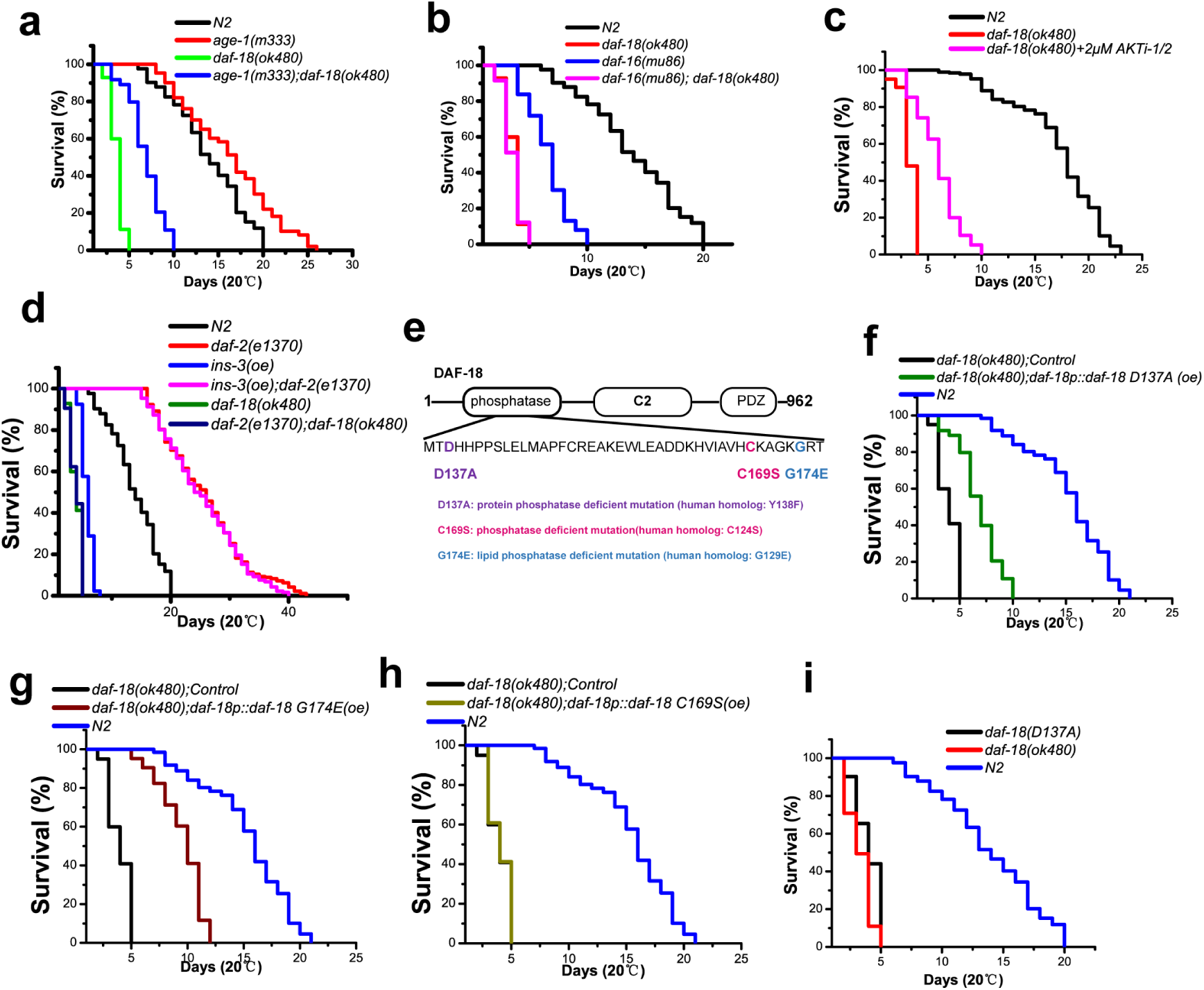
Both the lipid and protein phosphatase activities of DAF-18 affect L1 arrest survival. (a) Disruption of *age-1* partially rescued the shortened survival of *daf-18(ok480)* L1-arrested worms. (b) Disrupting *daf-18* reduced the survival of *daf-16* mutants even further. (c) The activation *of daf-16* failed to fully rescue the shortened survival of *daf-18(ok480)* L1-arrested worms. (d) Unlike *ins-3* worms, disruption of *daf-2* failed to rescue the short survival of *daf-18* worms. (e) Schematic of DAF-18 phosphatase and the mutations introduced in the experiments. Expression plasmids containing mutant sequences were used to generate the specific proteins. (f) The lipid phosphatase activity of DAF-18 partially rescued the survival of *daf-18(ok480)* L1-arrested worms. (g) The protein phosphatase activity of DAF-18 partially rescued the survival of *daf-18(ok480)* L1-arrested worms. (h) Overexpression of *daf-18* without phosphatase activity failed to rescue the shortened survival of *daf-18(ok480)* L1-arrested worms. (i) The effect of CRISPR/Cas9-modified DAF-18 lacking protein phosphatase activity on survival during L1 arrest. Each set of survival experiments was independently repeated at least three times. The mean survival rates were calculated using the Kaplan‒Meier method, and P values were determined by using the log-rank test. All the survival data are summarized in Table S1.

### linc-133 gain-of-function mutation rescues the shortened survival caused by daf-18 loss

To identify the gene mutations that can fully rescue the shortened survival of *daf-18* L1-arrested worms, a forward genetic screen was performed by using genome-wide mutagenesis with ethyl methanesulfonate (EMS) ^15,23^. Thousands of F3 EMS-mutagenized animals were screened, and two F3 progeny of EMS-treated *daf-18* worms were found to have normal L1 arrest survival (Fig. 2a). A single-nucleotide polymorphism in *linc-133,* which we termed *aqz1*, was identified in these mutants by using the Sibling Subtraction Method ^24^. *linc-133(aqz1)* fully recovered the survival of *daf-18* L1-arrested worms to that of the N2 worms (Fig. 2b). *linc-133* is predicted to be a non-coding RNA with no obvious role described so far. To further address the function of this new allele of *linc-133*, we made three RNAi versions to silence the expression of wild-type *linc-133* (Fig. 2c). We found that all three RNAi plasmids failed to rescue the survival of L1 arrest *daf-18* worms (Fig. 2d). Then, we used CRISPR/Cas9 to delete *linc-133* (termed *aqz2*) and to make the *aqz1* mutation (termed *aqz3)* (Fig. 2e). We found that the deletion version of *linc-133(aqz2)* failed to rescue the survival of L1 arrest *daf-18* worms (Fig. 2f). We also overexpressed wild-type *linc-133* in *daf-18* worms, but *linc-133(oe)* had no obvious effect on the survival of L1-arrested worms (Fig. 2g). However, *linc-133(aqz3)*, the CRISPR/Cas9 version of *linc-133(aqz1)*, fully rescued the survival of L1-arrested *daf-18(ok480)* worms (Fig. 2h). These results suggest that the *aqz1/3* alleles of *linc-133* may be gain-of-function mutations.

**Figure 2.**
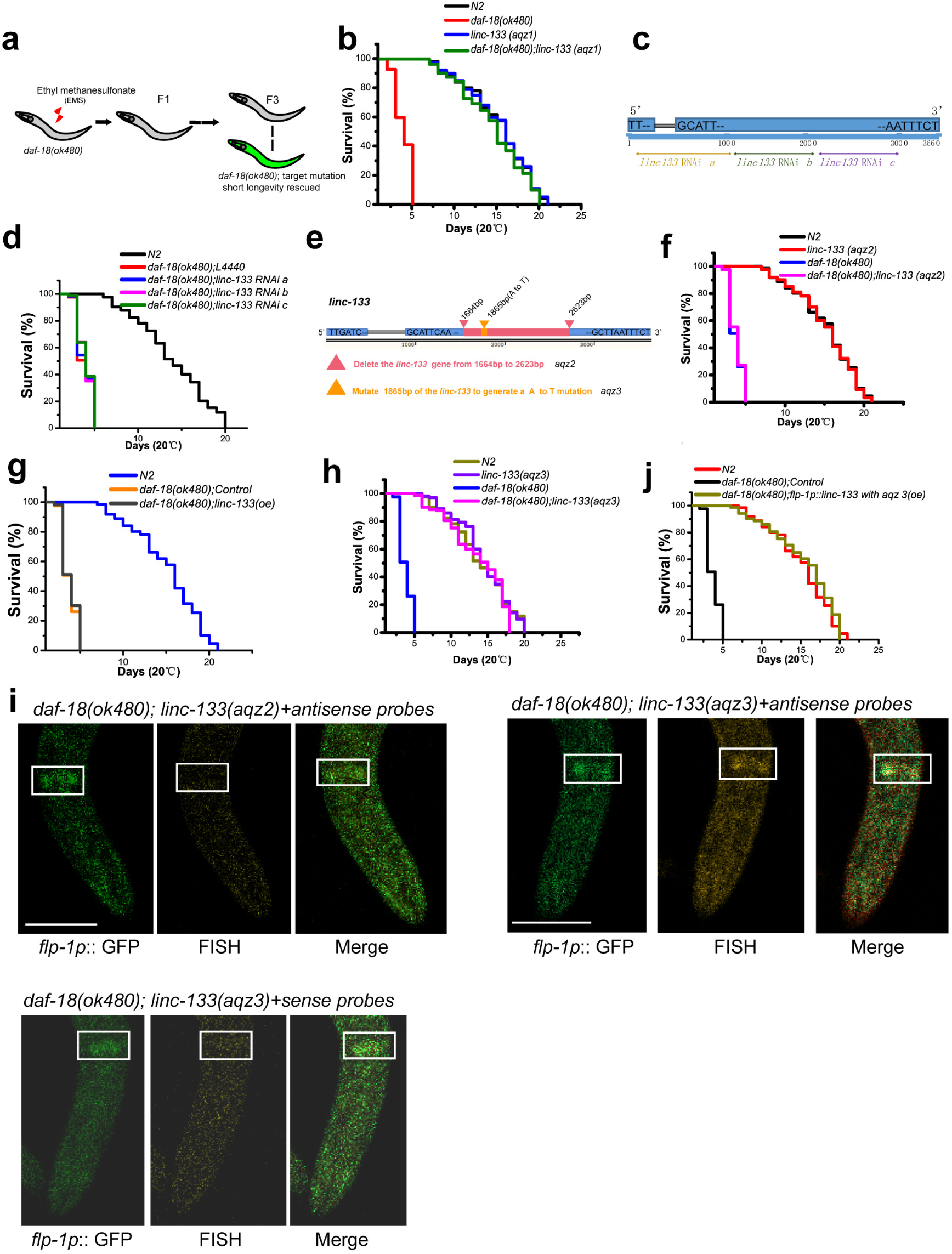
*linc-133* gain of function rescues the shortened survival of *daf-18(ok480)* L1-arrested worms. (a) EMS screening for suppressors of *daf-18*. *daf-18(ok480)* worms were used to perform the screening. (b) A mutation termed *aqz1* was identified to almost fully rescue the shortened survival of *daf-18(ok480)* L1-arrested worms. (c-d) Three different *linc-133* knockdown RNAi constructs were generated and tested for their effects on the survival of *daf-18(ok480)* L1-arrested worms. (e) CRISPR‒Cas9 engineered endogenous alleles included *aqz2* (*linc-133* deletion) and *aqz3* (CRISPR‒ Cas9 version of *aqz1*). (f) Deletion of *linc-133* failed to rescue the shortened survival of *daf-18(ok480)* L1-arrested worms. (g) Overexpression of *linc-133* failed to rescue the shortened survival of *daf-18(ok480)* L1-arrested worms. (h) The mutation *aqz3* (CRISPR‒Cas9 version of *aqz1*) fully rescued the shortened survival of *daf-18(ok480)* L1-arrested worms. Each set of survival experiments was independently repeated at least three times. The mean survival rates were calculated using the Kaplan‒Meier method, and P values were determined by using the log-rank test. All the survival data are summarized in Table S1. (i) *linc-133* RNA enrichment in AVK cells was tested by using *in situ* hybridization in whole-mount worms. The antisense probes were used to test *lin-133*, sense probes as negative control. (j) Overexpression of *linc-133 aqz3* driven by the *flp-1* promoter could extend the survival of *daf-18(ok480)* worms during L1 arrest. Scale bar: 25 µm. FISH: Fluorescence *in situ* hybridization, pale red. DAPI: 4’,6-Diamidino-2-phenylindole fluorescent staining.

We next wanted to know the location of its potential function inside the cells. Forty-eight short DNA nucleotide primers were used for RNA *in situ* hybridization to localize the mutated *linc-133* ^25,26^. *linc-133* was reported to be enriched in AVK cells, and our results confirmed that AVKs have high levels of mutated *linc-133* (Fig. 2i) and that overexpression of *linc-133* with *aqz3* mutation in AVK cells can significantly rescue the survival of L1 arrest *daf-18* worms (Fig. 2j), suggesting that *linc-133* gain of function may play a cell nonautonomous role. However, the overall signal (outside AVK cells) appears much higher in *linc-133 (aqz3)* mutants compared to *linc-133 (aqz2)* null mutants suggesting that *linc-133* may also be expressed outside AVK cells. In total, these results suggest that both lipid and protein phosphatase activities can be compensated by *linc-133* gain-of -function mutation and that *linc-133* is a common effector for both DAF-18 activities.

### linc-133 gain-of-function obstructs DAF-16 translocation from the nucleus to the cytoplasm

*linc-133* RNAi could abolish the survival extension of *linc-133(aqz3)* (Fig. 3a). To identify the molecular mechanism by which the *aqz3* allele of *linc-133* regulates survival, we performed RNA sequencing (RNA-seq) to compare the gene expression profile changed by *linc-133(aqz3)* mutation (Fig. 3b). The Kyoto Encyclopedia of Genes and Genomes (KEGG) pathway enrichment results showed that one set of genes with up-regulated expression in *linc-133(aqz3)* worms was enriched in the DAF-16-regulated survival pathway (Fig. 3c). We proposed that *linc-133(aqz3)* might regulate the function of the transcription factor DAF-16; however, disruption of *daf-16* only partial reduced the survival of *linc-133(aqz3)* worms during L1 arrest (Fig. 3d). The double mutants of *daf-16(mu86); linc-133(aqz3)* live significantly longer than *daf-16* worms; however, disruption of *daf-16* failed to further shorten the survival of *daf-18(ok480);linc-133(aqz3)* (Fig. 3d). These results suggested that DAF-16 is one of the targets of *linc-133(aqz3),* and *linc-133(aqz3)* can compensate the loss of DAF-16 through other effects. This is consistent with the previous report that *daf-18* and *daf-16* do not have additive effects for L1 starvation survival ^27^. To confirm whether the *linc-133* gain-of-function mutant regulates the function of DAF-16, we performed a DAF-16 nuclear localization experiment in living worms. Our results showed that the localization of DAF-16::GFP was enhanced by the *aqz3* allele of *linc-133* (Fig. 3e). Moreover, the typical DAF-16 target gene *sod-3* was up-regulated in living *linc-133(aqz3)* L1-arrested worms (Fig. 3f-g).

**Figure 3.**
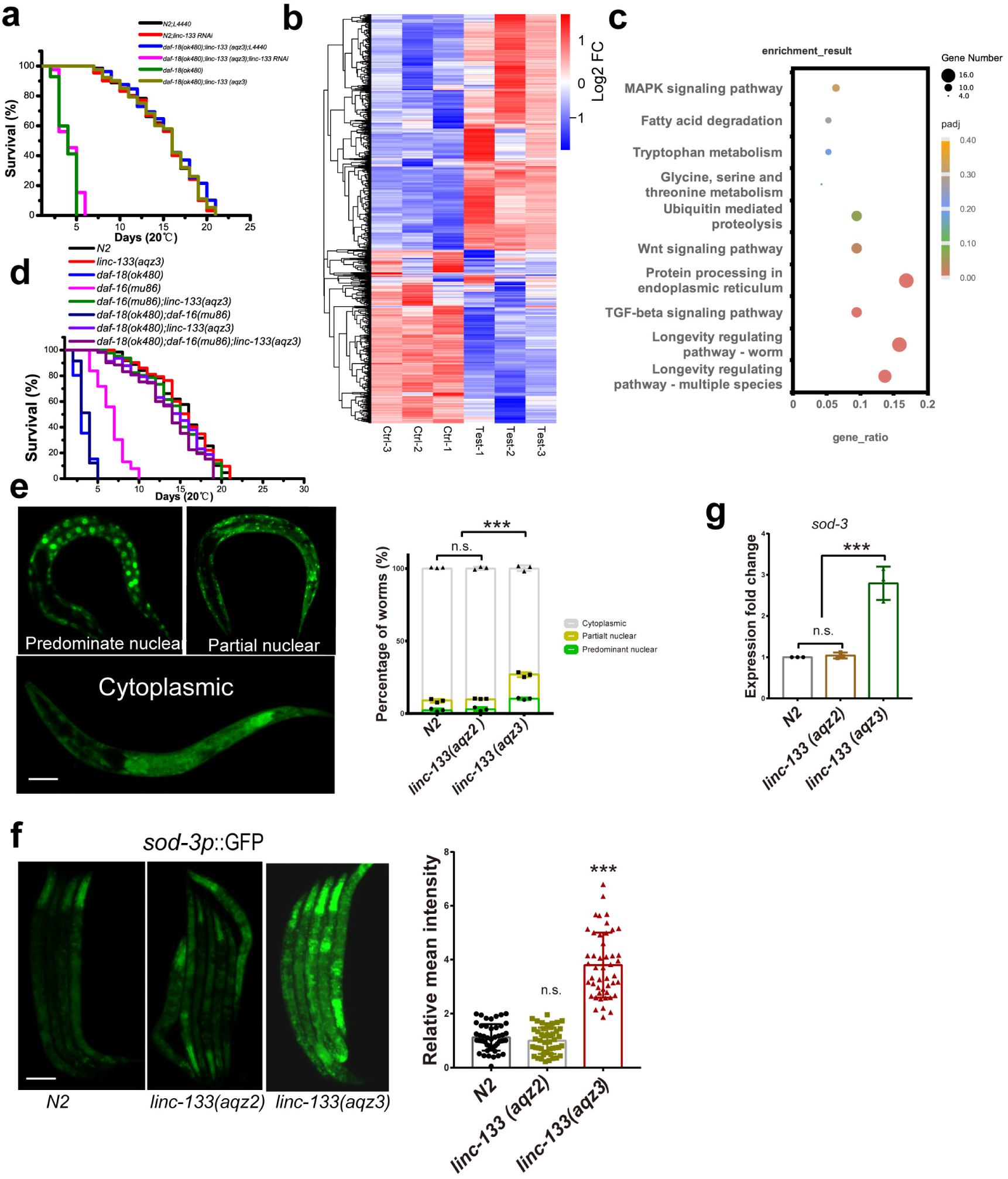
*linc-133* gain-of-function partially regulates DAF-16. (a) *linc-133* RNAi abolished the extended survival. (b) Comparison of the gene expression in gain-of-function *linc-133* worms and wild-type worms. Genes with expression changes of more than twofold according to RNA-seq were selected. Ctrl: *daf-18(ok480)*, Test: *daf-18(ok480); linc-133(aqz3)*. (c) KEGG pathway enrichment of genes upregulated in gain-of-function of *linc-133* worms. (d) *linc-133* gain-of-function regulate survival partially dependent on DAF-16 during L1 arrest. (e) Measurements of DAF-16 nuclear localization. n.s.: no significant difference. ***: p<0.001. Scale bar: 50 µm. (f) *sod-3p::GFP* was used to test the expression of DAF-16 target genes. Scale bar: 50 µm. n.s.: no significant difference. ***: p<0.001. (g) The expression of *sod-3* was tested by using real time PCR.

Considering that DAF-16 will be translocated to the cytoplasm from nuclei when worms lose *daf-18*, we proposed that the *linc-133(aqz3)* may obstruct DAF-16 translocation. Then, we heat shocked the *daf-18* and *daf-18;linc-133* double mutant worms at 34℃ to make DAF-16 localized in nuclei (Fig. 4a). When the relocation of nuclear accumulated DAF-16 to the cytoplasm was induced by low-temperature (20℃) treatment or AKT activation (treated with AKT activator SC79), *linc-133(aqz3)* made the *daf-18* worms have more cells with nuclear localized DAF-16 than *linc-133* deletion worms (Fig. 4b-c). Next, we asked how *linc-133(aqz3)* obstructs translocation out of the nucleus. We speculated that the *linc-133* gain-of-function mutant may bind to the cofactors of DAF-16, so we performed RNA pulldown with *linc-133* nucleic acid probes. Results showed that a specific protein interacted with *linc-133* probes in *linc-133 (aqz3)*, and *daf-18(ok480);linc-133(aqz3)*, but not in *daf-18(ok480)* worms (Fig. 4d). Mass spectrometry identified the protein is 14-3-3. The 14-3-3 protein FTT-2 forms a complex with DAF-16 and prevents DAF-16 from translocation ^28,29^. Our result suggests that the wild-type *linc-133* can interact with 14-3-3 protein, but this interaction is lost when *daf-18* is dysfunctional.

**Figure 4.**
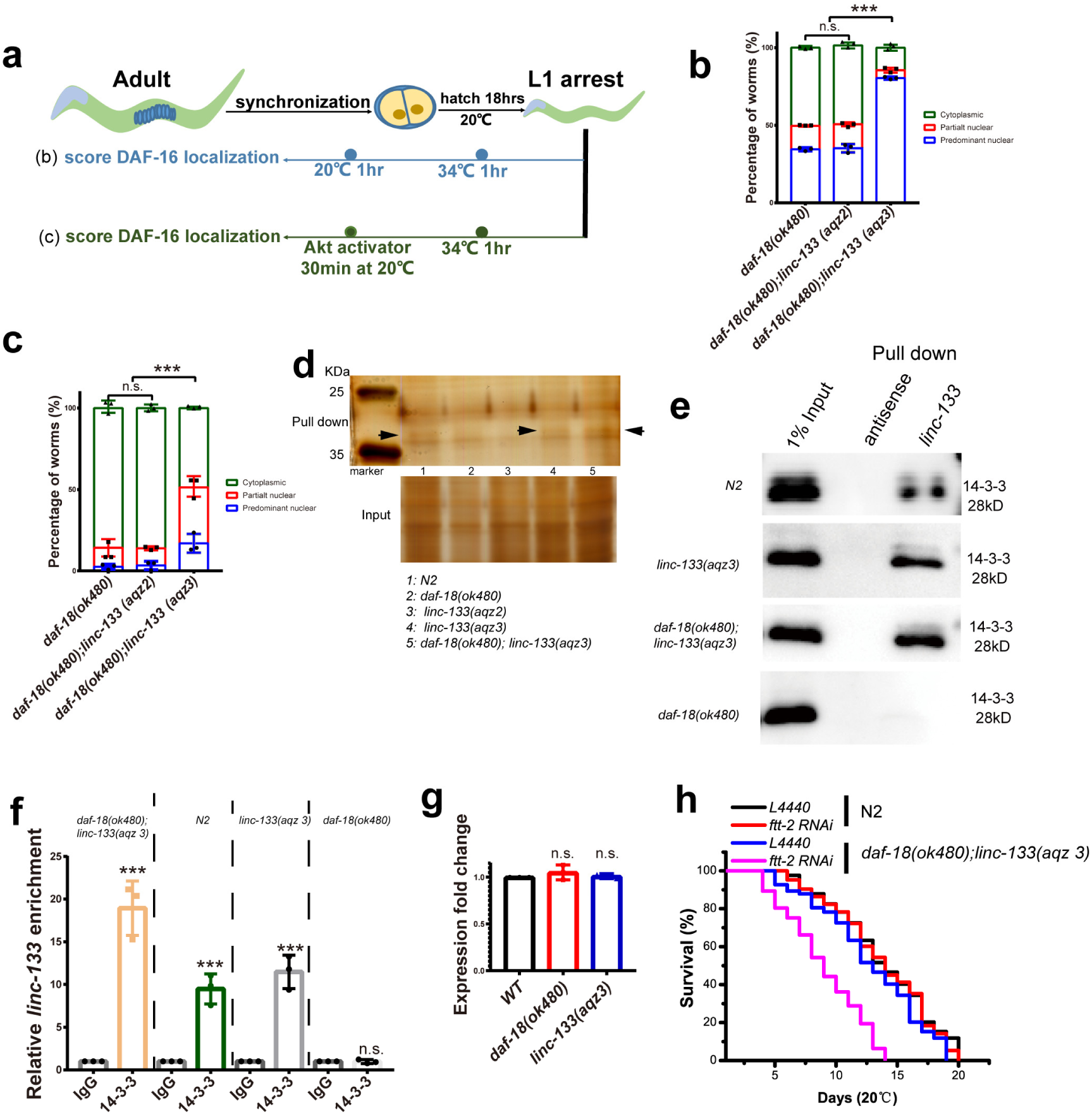
*linc-133* gain of function affects the translocation of DAF-16. (a) Schematic of the experiments performed to test DAF-16 translocation from the nucleus to the cytoplasm. (b-c) Measurements of DAF-16 nuclear localization. n.s.: no significant difference. ***: p<0.001. (d) *linc-133* probes were used to perform RNA pulldown in N2, *daf-18(ok480), linc-133(aqz2), linc-133(aqz3), daf-18(ok480);linc-133(aqz3)* worms. Mass spectrometry identified the 14-3-3 proteins (The complete list of proteins identified by mass spectrometry is included in Supplementary data). (e) The *linc-133* probes were used to perform RNA pulldown, and the specific 14-3-3 antibody was used to do the blotting. (f) The 14-3-3 antibody was used to co-immunoprecipitated 14-3-3 proteins and *linc-133* RNA. The RIP confirmed the *linc-133 aqz3* can be immunoprecipitated together with 14-3-3 antibody. (g) The expression of *linc-133* was not affected by *daf-18* and *linc-133 aqz3*. (h) Knocking down 14-3-3 coding genes affected the survival of *daf-18(ok480);linc-133(aqz3)* worms. The mean survival rates were calculated using the Kaplan‒Meier method, and P values were determined by using the log-rank test. All the survival data are summarized in Table S1.

Interestingly, the interaction between *linc-133* and 14-3-3 protein is enhanced by *aqz3* mutation in N2 or *daf-18* worms. *linc-133* probes were used to perform RNA pulldown and the specific 14-3-3 antibody staining further confirmed that the mutated *linc-133 aqz3* physically connects with 14-3-3 proteins (Fig. 4e). Moreover, we also used RNA binding protein immunoprecipitation (RIP) to test whether the mutated *linc-133 aqz3* can be immunoprecipitated together with 14-3-3 proteins. The real-time PCR showed that the *linc-133* is enriched by 14-3-3 antibody in *linc-133 aqz3* worms (Fig. 4f). The expression of mutated *linc-133* is not changed in *daf-18(ok480)* and *linc-133(aqz3)* worms (Fig. 4g), suggesting the interaction between mutated *linc-133* and 14-3-3 protein is not attributed to expression dosage. Further, knocking down *ftt-2* reduced the survival of *daf-18(ok480);linc-133(aqz3)* during L1 arrest (Fig. 4h). These results suggest that the *linc-133* gain-of-function mutant physically interacts with 14-3-3 to block DAF-16 from exiting the nucleus in *daf-18* worms.

### linc-133 gain of function works through HSF-1

As DAF-18 protein phosphatase activity plays a pivotal role in supporting survival during L1 arrest, we next asked how *linc-133(aqz3)* compensates for DAF-18 protein phosphatase defects. According to our RNA-seq analysis, the most significant gene expression changes in *linc-133(aqz3)* worms were enriched in protein degradation in the endoplasmic reticulum pathway (Fig. 3c). The enriched genes were up-regulated in *linc-133(aqz3)* worms (Fig. 5a), and their expression was confirmed by real-time PCR (Fig. 5b). These genes were down-regulated in *daf-18(ok480)* (Fig. 5c) and *daf-18(D137A)* worms (Fig. 5d), but up-regulated by the mutation of *aqz3* in *daf-18 (ok480);linc-133(aq3)* (Fig. 5e) and *daf-18 (D137A);linc-133(aqz3)* worms (Fig. 5f). Most of these genes are heat shock genes which regulated by the transcription factor HSF-1 ^30^, so next we test whether the function of *linc-133(aqz3)* is related to HSF-1. Our results showed that disruption of *hsf-1* significantly reduced the survival extension of *linc-133(aqz3)* (Fig. 5g), suggesting HSF-1 is the main target of *aqz3* to regulate survival of L1 arrest worms. Then, we tested whether the HSF-1 is main target of *linc-133(aqz3)* in *daf-18* worms. Our results showed the rescuing functions of *linc-133(aqz3)* on the survival of *daf-18* worms were significantly decreased by disruption of *hsf-1* (Fig. 5h). However, disruption of *hsf-1* failed to totally abolish the survival extension of *linc-133(aqz3)*, also suggesting that HSF-1 is not the only target of *linc-133(aqz3).* Interestingly, the survival of *daf-18 (D137A); linc-133(aqz3)* was abolished to that of *daf-18 (D137A)* worms (Fig. 5i). These results suggest that the DAF-18 protein phosphatase activity may mainly regulate HSF-1. We knocked down all these enriched heat shock genes regulated by HSF-1, in *daf-18(ok480);linc-133(aqz3)* and *daf-18 (D137A); linc-133(aq 3)* worms and found that knocking down *hsp-1* significantly reduced the survival of *daf-18(ok480);linc-133(aqz3)* (Fig. 5j) and *daf-18 (D137A); linc-133(aqz3)* (Fig. 5k) worms during L1 arrest. Taking together, the results suggest that the dysfunction of protein phosphatase activity of DAF-18 is rescued by *linc-133(aqz3)* through regulating HSF-1 to maintain the survival of *daf-18* worms.

**Figure 5.**
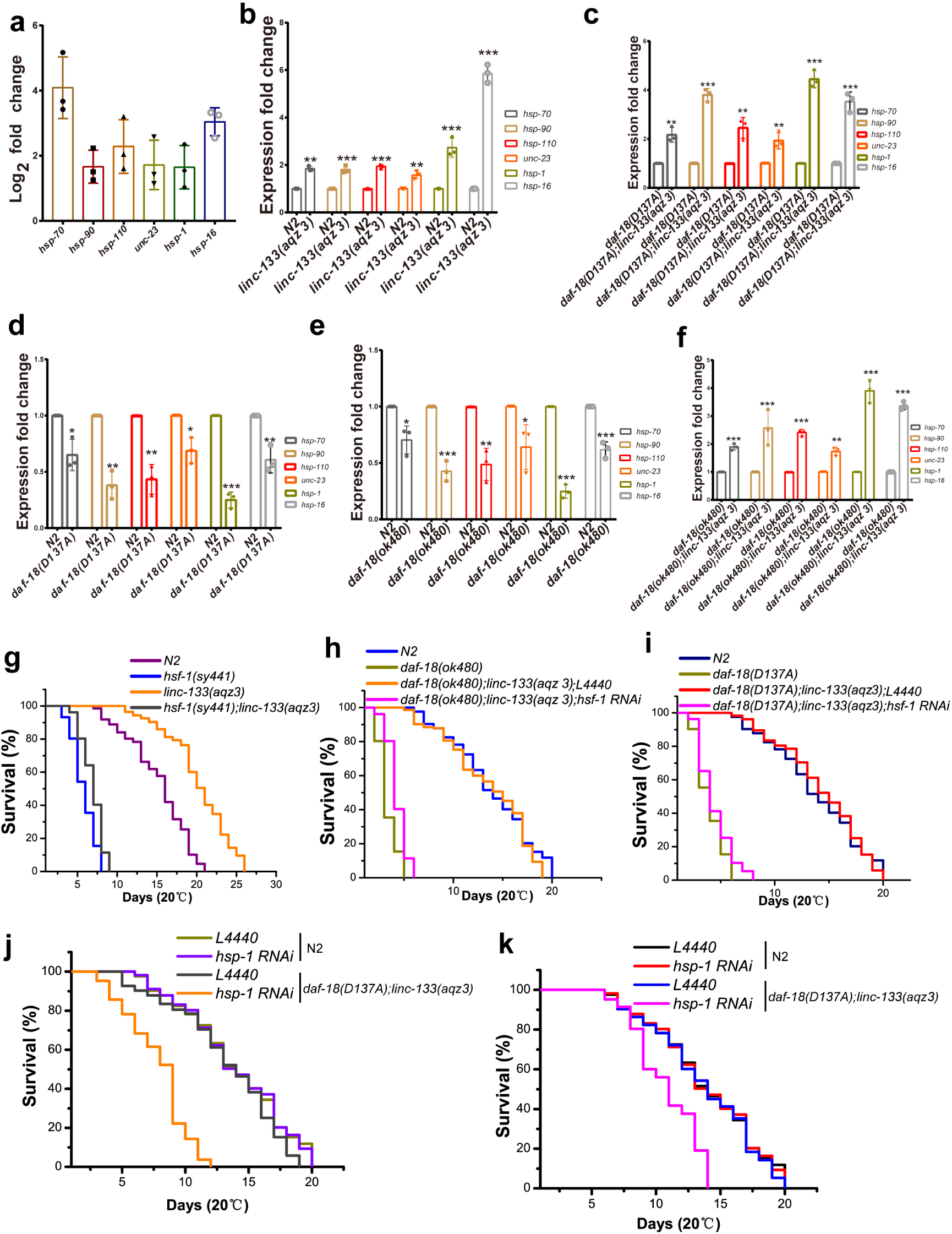
linc-133 gain-of-function regulates HSF-1 to support the survival when the protein phosphatase activity of DAF-18 is dysfunctional. (a) Average changes in gene expression of chaperones according to RNA-seq. (b-f) The fold changes in gene expression were tested by using real-time PCR. *: p<0.05, **: p<0.01, ***: p<0.001. (g) The effects of *hsf-1* on survival of *linc-133 (aqz3)* worms during L1 arrest. (h) The effects of *hsf-1* on survival of *daf-18 (ok480)*; *linc-133 (aqz3)* worms during L1 arrest. (i) The effects of *hsf-1* on survival of *daf-18 (D137A)*; *linc-133 (aqz3)* worms during L1 arrest. (j) Knocking down *hsp-1*, a target gene of HSF-1, decreased the survival of *daf-18 (ok480)*; *linc-133 (aqz3)*. (k) Knocking down *hsp-1*decreased the survival of *daf-18 (D137A)*; *linc-133 (aqz3)* worms. Each set of survival experiments was independently repeated at least three times. The mean survival rates were calculated using the Kaplan‒Meier method, and P values were determined by using the log-rank test. All the survival data are summarized in Table S1.

### linc-133 gain of function enhances protein degradation

According to the RNA seq results, most of the genes, enriched in the top four pathways (Fig. 3c), actually encode the ubiquitous molecular chaperones HSP70, HSP90 and nucleotide exchange factor (NEF), which participate broadly in preventing protein aggregation and promoting the refolding of misfolded denatured proteins, solubilizing aggregated proteins and cooperate with cellular degradation machineries to clear protein aggregates ^31^. When aggregated proteins are produced, these chaperons promote the degradation of aberrant proteins by the ubiquitin proteasome system to help maintain protein homeostasis and ensure survival (Fig. 6a) ^31,32^. We speculated that the pressure of *daf-18* loss may cause the cells to produce large amounts of aggregated proteins, and *linc-133(aqz3)* induces HSP chaperones to degrade these protein aggregates. To test this speculation, we used the protein aggregation marker *Pnmy-2*NMY-2::GFP (aggregation-prone protein) to measure the level of protein aggregation ^33^. Our results showed that loss of *daf-18* or loss of the protein phosphatase activity of DAF-18 caused the worms to produce high level aggregated proteins during L1 arrest, and *linc-133(aqz3)* significantly decreased the levels of protein aggregations (Fig. 6b). The results also showed that a mutant *daf-18* rescue plasmid lacking the protein phosphatase activity failed to reduce the level of aggregated proteins (Fig. 6c). These results suggest that the protein phosphatase activity of DAF-18 controls protein aggregation. As our results showed that *linc-133(aqz3)* can rescue the dysfunction of protein phosphatase activity of DAF-18 through regulating HSF-1, as a transcription factor, which may play a key role in controlling unfolded proteins degradation. Indeed, knocking down *hsf-1* significantly increased the level of aggregated proteins in *daf-18(ok480);linc-133(aqz3)* (Fig. 6d) and *daf-18(D137A);linc-133(aqz3)* (Fig. 6e) L1-arrested worms. Next, we also knocked down *hsp-1*, the target gene of HSF-1, in these worms, we found that the levels of aggregated proteins in *daf-18(ok480);linc-133(aqz3)* (Fig. 6f) and *daf-18(D137A);linc-133(aqz 3)* (Fig. 6g) were also significantly increased.

**Figure 6.**
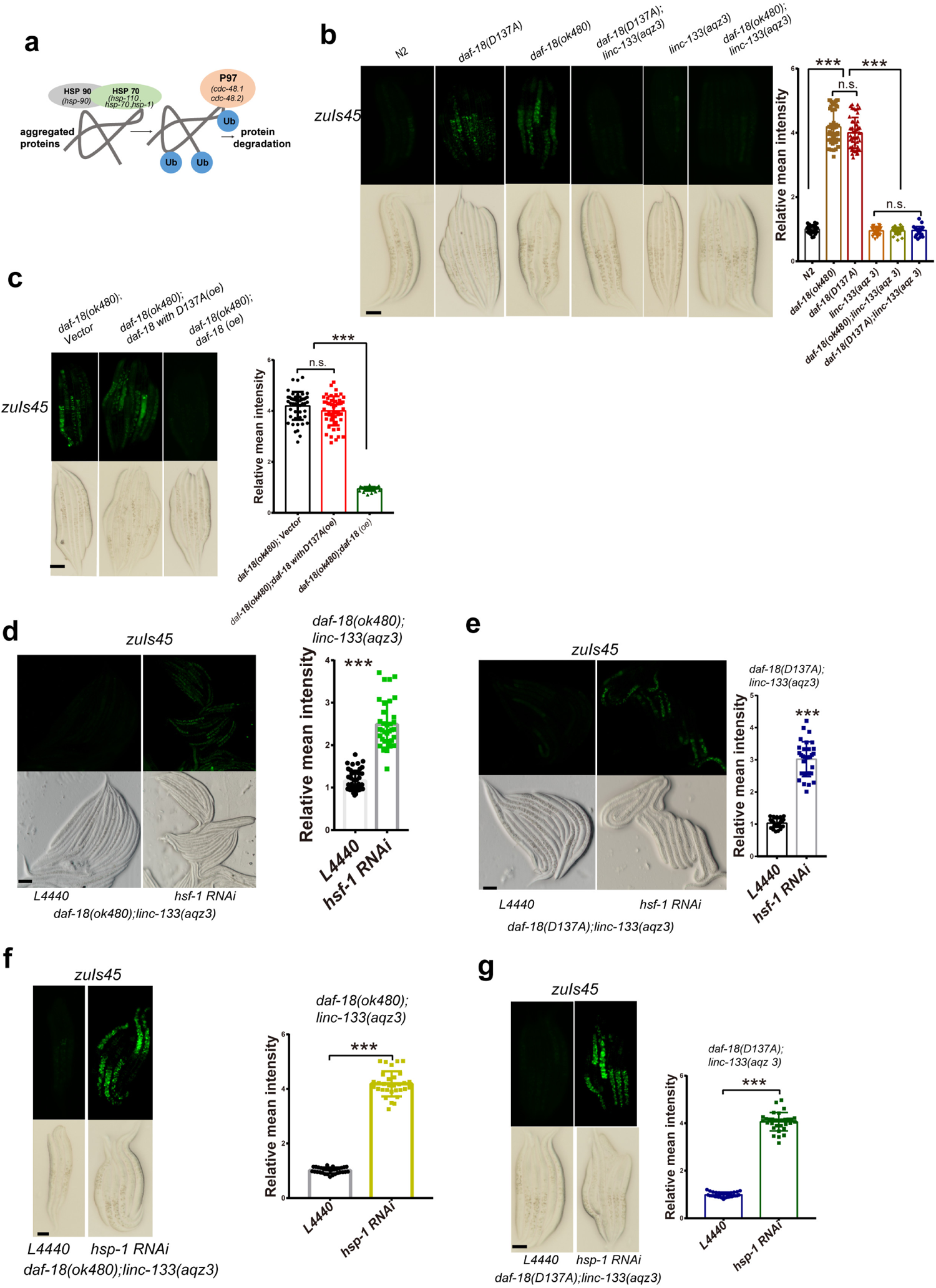
*linc-133* gain of function decreases the protein aggregation caused by loss of protein phosphatase activity of DAF-18. (a) Aggregated proteins selected and degraded in cytosol. (b) The protein aggregation affected by *daf-18(D137A)* and *linc-133* gain of function were identified by using the *zuIs45* (*Pnmy-2*NMY-2::GFP) marker. (c) The protein aggregation rescuing tests were identified by using the *zuIs45* (*Pnmy-2*NMY-2::GFP) marker. (d-e) The protein aggregation affected by *hsf-1* in *daf-18(ok480);linc-133(aqz3)* and *daf-18(D137A);linc-133(aqz3)* worms was identified by using the *zuIs45* (*Pnmy-2*NMY-2::GFP) marker. (f-g) The protein aggregation affected by *hsp-1* in *daf-18(ok480);linc-133(aqz3)* and *daf-18(D137A);linc-133(aqz3)* worms were identified by using the *zuIs45* (*Pnmy-2*NMY-2::GFP) marker. ***: p<0.001. Scale bar: 50 µm.

The aggregated proteins in cytoplasm can be selected, ubiquitinated and degraded by the ubiquitin proteasome system ^34–37^ (Fig. 6a). To test whether the ability of protein ubiquitination is affecting the survival of *linc-133*(*aqz3*) worms, we knocked down all the genes related to ubiquitination in ERAD, and found that knocking down the P97 coding genes *cdc-48.1* or *cdc-48.2* shortened the survival of *daf-18(ok480);linc-133(aqz3)* (Fig. 7a) and *daf-18(D137A);linc-133(aqz3)* (Fig. 7b) worms during L1 arrest. These results show that *linc-133(aqz3)* works through protein degradation to support survival during L1 arrest. To further confirm this model, we tested whether the protein ubiquitination levels are changed in *daf-18* and *linc-133* worms. Recent study has shown that short-lived worms exhibit higher protein aggregation and lower ubiquitin protein levels. In contrast, long-lived worms have elevated levels of ubiquitinated proteins, which are degraded by the proteasome to reduce the accumulation of aggregated and unfolded proteins ^38,39^. Our results showed that *daf-18(ok480)* and *daf-18(D137A)* (Fig. 7c) mutants had lower levels of K48-linked ubiquitinated proteins, and *linc-133(aqz3)* induced high levels of K48-linked ubiquitinated proteins (Fig. 7d). When we knocked down the P97 ubiquitin-selective chaperone coding gene *cdc-48.2*, the K48-linked ubiquitinated protein levels were decreased (Fig. 7e). Together, these results suggest that the loss of *daf-18* leads to higher levels of aggregated proteins and that the protein phosphatase activity of DAF-18 may play a pivotal role in this process. *linc-133(aqz3)* can induce HSF-1- and P97-related protein ubiquitination degradation to maintain protein homeostasis.

**Figure 7.**
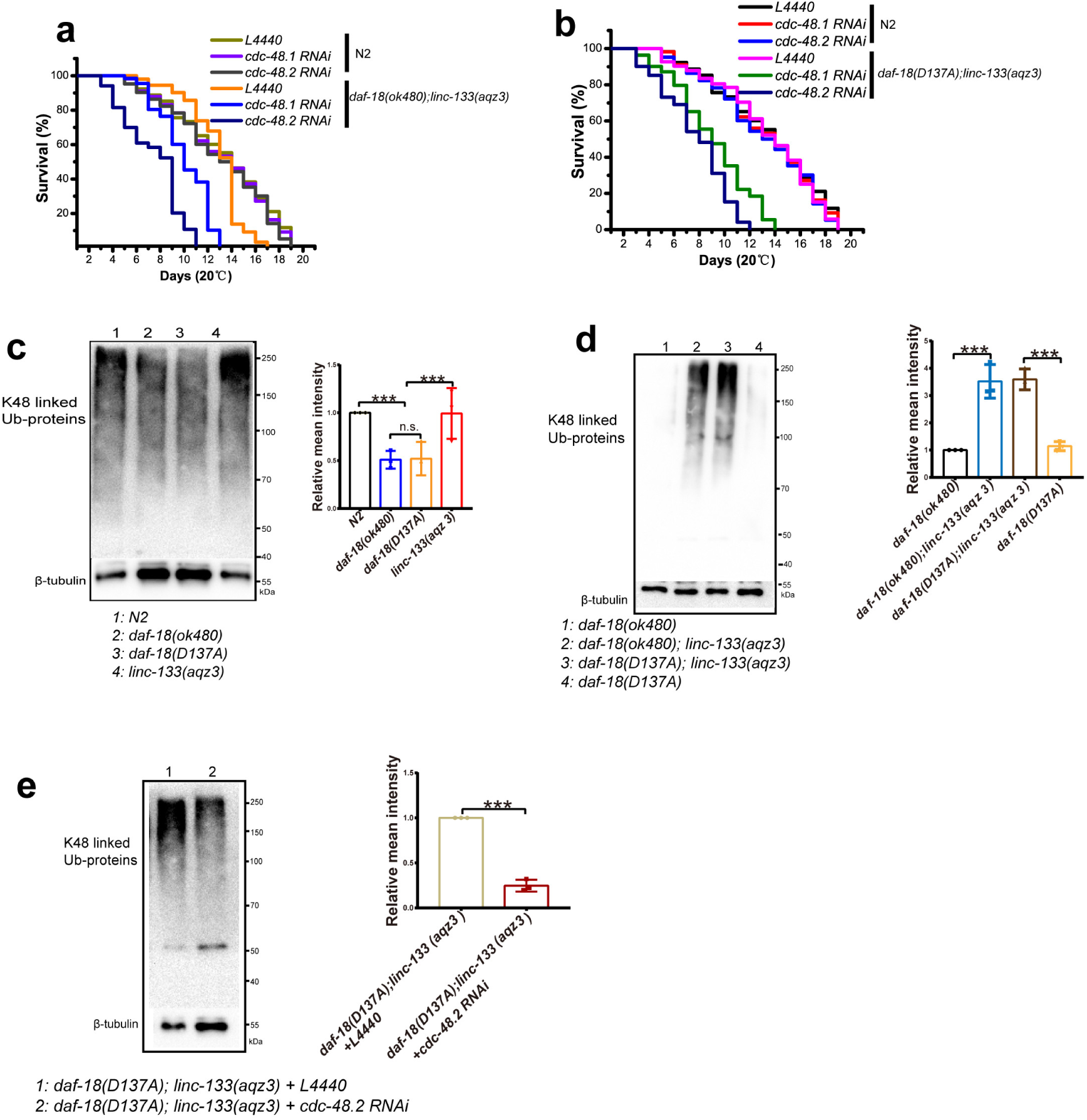
*linc-133* gain of function maintains protein homeostasis to support survival. (a-b) Disrupting P97 coding genes reduced the survival of *daf-18(ok480);linc-133(aqz3)* and *daf-18(D137A);linc-133(aqz3)* worms during L1 arrest. Each set of survival experiments was independently repeated at least three times. The mean survival rates were calculated using the Kaplan‒Meier method, and P values were determined by using the log-rank test. All the survival data are summarized in Table S1. (c) K48-linked protein ubiquitination in N2, *daf-18(ok480), daf-18(D137A),* and *linc-133(aqz3)* worms during L1 arrest. (d) K48-linked protein ubiquitination in *daf-18(ok480);linc-133(aqz3)* and *daf-18(D137A);linc-133(aqz3)* worms during L1 arrest. Each immunoblot is representative of three independent experiments. (e) K48-linked protein ubiquitination in *cdc-48.2* knockdown worms during L1 arrest. Columns show the average fold changes in K48-linked protein ubiquitination in three independent experiments. n.s.: no significant difference. ***: p<0.001. Scale bar: 50 µm.

## Discussion

Loss of *daf-18* leads to two prominent phenotypes in L1 arrested worms: cell proliferation and survival decrease. Previous research has extensively studied cell divisions, including P, V, Q, and germ cell divisions, in *daf-18* worms during L1 arrest, and many genes and signaling pathways such as TOR, TGF-beta, IIS, and MAPK have been implicated ^4,5,8,40^. However, none of these genes were capable of fully rescuing the short survival of *daf-18* L1 arrested worms (Fig. S1), despite some studies showing that IIS, TGF-beta have some functions on the survival of L1 arrested worms ^2,3,5,7,9,41,42^. We used a traditional forward genetic screening method to identify a new gene mutation that can compensate for the loss of *daf-18/PTEN* (Fig. 8). In addition to the lipid phosphorylation of DAF-18 and related PI3K signaling, the protein phosphatase activity of DAF-18 also plays important roles in controlling survival. These findings are consistent with recent reports that protein phosphatase activity of DAF-18/PTEN has roles in controlling oncogenic signaling independently of PI3K-AKT^21,43,44^. We demonstrated that the protein phosphatase activity of DAF-18 is critical for the regulation of aggregated protein and protein homeostasis. *linc-133 aqz3* can compensate for this by regulating the gene expression of the ERAD pathway, specifically by evoking the HSF-1 pathway to regulate heat shock chaperones that can select and identify aggregated proteins in the cytoplasm. These aggregated proteins can then be ubiquitinated and degraded, thereby maintaining protein homeostasis and supporting the survival of *daf-18* mutants during L1 arrest. This is consistent with previous reports that short-lived worms, such as *daf-18* mutants, have higher levels of unfolded and aggregated proteins and lower levels of ubiquitinated proteins, indicating a weakened ubiquitin degradation system ^38^. *daf-18* worms are more susceptible to heat stress and exhibit reduced mobility, likely due to the accumulation of aggregated proteins ^45^. Our results also showed that the *linc-133(aqz3)* increased resistance to heat stress and improved mobility of *daf-18* worms (Fig. S2). This further supports the conclusion of the study. We speculate that the function of HSF-1 can be independently evoked by *linc-133(aqz3)*, despite the fact that the IIS pathway can also regulate HSF-1. However, we cannot exclude the possibility of inter-regulation between HSF-1 and the IIS pathway in *linc-133(aqz3)* worms based on our results. It is possible that *linc-133(aqz3)* regulates HSF-1 independently of IIS, as previous studies have reported that HSF-1 is regulated by IIS^46^, but still has functions independent of DAF-16 ^47–50^.

**Figure 8.**
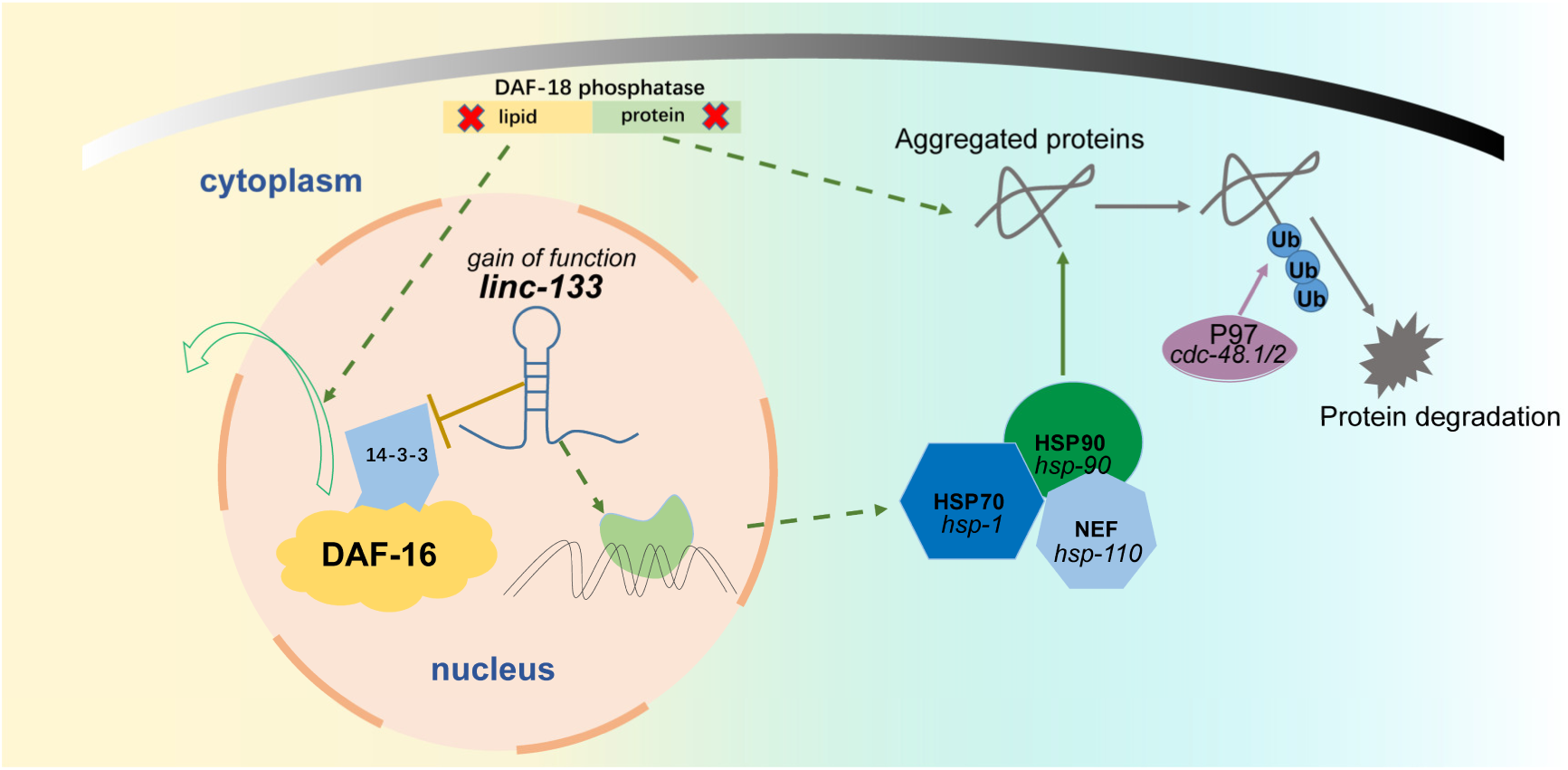
The working model of *linc-133* gain-of-function in *daf-18*-deficient L1-arrested worms. Defects in lipid phosphatase activity can cause translocation of the transcription factor DAF-16 from the nucleus to the cytoplasm, and *linc-133* gain-of-function may obstruct this translocation. The gain of function of *linc-133* may also be involved in other AGE-1/PI3K-regulated biological processes through regulating HSF-1. Protein phosphatase activity plays a pivotal role in controlling L1 arrest survival. Loss of DAF-18 protein phosphatase activity causes the production of high levels of aggregated proteins in L1-arrested worms. The gain-of-function of *linc-133* can induce HSP chaperones to select and process the aggregated proteins for degradation through ubiquitination modification.

The targets and mechanisms of *linc-133* gain-of-function mutations that we identified in the L1 arrest stage may also be applicable to other life stages in worms. Considering that the human ortholog of *daf-18* is *PTEN*, a critical tumor suppressor gene, our findings may have implications for the study of PTEN-related diseases. It is worth exploring whether there are additional potential orthologs of *linc-133* or alternative genes that have this function in humans.

## Methods and materials

### *C. elegans* strains and culture

*C. elegans* strains were maintained using standard methods ^15^. All strains were cultured and maintained at 20 °C unless noted otherwise. All hybrid strains were generated in our laboratory and are described in the Supplementary Materials and Methods. All *C. elegans* were L1-arrested hermaphrodites. Double-mutant strains were created by crossing males of one strain with hermaphrodites of the other strain. Double mutants were checked by extracting their DNA, amplifying a genomic fragment flanking the mutation site by PCR, and Sanger sequencing of the PCR product. The strains used in this study are as follows: N2, RB712: *daf-18(ok480)*, TU3401: *sid-1(pk3321)*; *uIs69*, VC222: *raga-1(ok386), F*DR722: *age-1(m333)/mnC1[dpy-10(e128)unc-52(e444)*], TJ356: *zIs356*, JJ1473: *zuIs45*, CF1553: *muIs84*, CF1038: *daf-16(mu86)*. CB1370: *daf-2 (31370)*, PS3551: *hsf-1(sy441),* VC546: *daf-4(ok827)*, VC1183: *sma-9(ok1628)*, VC1270: *lin-31(gk569)*, CB1372: *daf-7(e1372)*, SD420: *mpk-1(ga119)/dpy-17(e164) unc-79(e1068)*, MU48: *lin-45(n2018)/dpy-20(e1282*), MT8666: *mek-2(n1989),* P*ins-3::gfp* (wwIs26), VC3044: *dbl-1(ok3749)*, ZG31: *hif-1(ia4)*, RB1206: *rsks-1(ok1255)*, AA1: *daf-12(rh257)*, MU1085 *bwIs2* [*flp-1::GFP + rol-6(su1006)*].

### *C. elegans* RNAi

RNAi feeding was conducted according to standard protocols ^51^. The RNAi bacteria were prepared and seeded onto NGM plates containing 1 mM isopropyl-B-D-thiogalactopyranoside (IPTG). Eggs were grown on NGM plates until the L4 stage and then transferred to RNAi plates. *C. elegans* strains were well fed before phenotypic analysis. RNAi bacteria containing the empty vector L4440 were used as controls. The RNAi constructs were obtained from the Horizon RNAi library (Horizon Discovery Ltd). To silence other designated target genes, the PCR products were cloned into the bacterial expression vector L4440 between opposing phage T7 polymerase promoter sites. The feeding vector was then transformed into *E. coli* HT115. The efficiency of RNAi was confirmed by using real time PCR. The primers used to construct these RNAi plasmids are presented in Table S2. All worms used to perform RNAi experiments contained the neuronal SID-1 receptor unless otherwise stated.

### Real-time PCR

L1-arrested worms sufficient to yield 50-100 µL of a tightly packed worm pellet were collected, and 1–1.5 µg/µL total RNA was extracted. Total RNA was isolated using RNAiso Plus (Takara) according to the manufacturer’s protocol, and cDNA was generated using a HiScript II 1st Strand cDNA Synthesis Kit (Vazyme). Real-time PCRs were run using Cham Q Universal SYBR qPCR Master Mix (Vazyme) on a Roche LightCycler 480Ⅱ. The relative expression levels of the genes were assessed using the 2^-ΔΔCt^ method and normalized to the expression of the *act-1* gene. P values were calculated using a two-tailed t test ^3,52^. The primers for real-time PCR are listed in Table S2.

### L1 arrest survival assay

Mixed-stage worms were collected and blenched in Eppendorf tubes (volume: 2 mL) to collect embryos. The embryos were resuspended and hatched in sterile M9 and incubated at 20 °C with low-speed rocking to enter L1 arrest. Two days later, a 20-50 µL aliquot with more than 50 L1-arrested worms was plated onto a 6 cm unseeded NGM plate. The numbers of live and dead worms were counted daily until all the larvae died. The percentages of live worms were calculated every day. The survival rates between strains were compared as described previously ^3,53^, and the mean survival rate was calculated by the Kaplan‒Meier method. The P values of the difference in overall survival rate were determined using the log-rank test.

### RNA sequencing

Wild-type N2 and *linc-133(aqz 3)* L1-arrested worms were collected. RNA-seq was completed by Novogene Corporation (http://www.novogene.com). RNA was prepared using Trizol reagent. Total RNA quality was measured using a NanoDrop ND-1000 (Agilent Technologies). Illumina TruSeq RNA Sample Prep Kit (Cat#FC-122-1001) was used with 1 µg of total RNA for the construction. RNA libraries were prepared for sequencing using standard Illumina protocols. Paired-end sequencing was performed on an Illumina NovaSeq 6000. Transcriptome alignment and quantification were performed using Hisat2 software. *C. elegans* genome version WBcel235 was used as the reference. KEGG pathway enrichment was performed using clusterProfiler software and the most up-to-date version of the KEGG database.

### Fluorescence *in situ* hybridization

For smFISH, we used custom Stellaris probes specific for *linc-133* lncRNA (48 probes) labeled with Quasar570 (excitation 548 nm, emission 566 nm). The probes were designed by Biosearch Technologies. The sequences of the new *linc-133* probes can be found in Table S2. Sample preparation and hybridization were performed in tubes using previously described protocols with some modifications ^25^. Images were recorded with a Leica TCS SP8 STED camera.

### GFP fluorescence assay

To measure GFP fluorescence in L1-arrested worms, the day one L1-arrested worms were transferred onto 2% agar pads. Images were taken under a fluorescence microscope (Nikon DS-RI2). For each strain, at least three independent replicate experiments were carried out with at least 30 worms of each strain captured in each experiment. The relative GFP intensity was analyzed and calculated by using ImageJ. The P values were determined by using a two-tailed t test.

### DAF-16::GFP localization assay

To test DAF-16 translocation, a previously described standard method was used ^52^. Worms were maintained at 20 °C, transferred onto 2% agar pads for observation of DAF-16::GFP nuclear localization status. To test DAF-16 translocation from the nucleus, day one L1-arrested worms were heat shocked at 34 °C for 1 h to allow DAF-16 translocation into the nucleus. Then, the worms were either kept at 20 °C or treated with an Akt activator (SC79, diluted in M9 buffer; working concentration: 10 mg/kg SC79) for another half hour before observation. At least 80 animals were observed per treatment, and each treatment included three independent repeats. The localization status of DAF-16::GFP was scored and assigned to three categories: predominantly nuclear, partially nuclear and cytoplasmic. The location of DAF-16::GFP was monitored using a fluorescence microscope system. ((Nikon DS-RI2)). Accumulation of fluorescent signal in nuclei was scored as described previously ^52^. P values were calculated using two-way ANOVA.

### Transgenic strains

The *linc-133* deletion and gain-of-function mutant worms were generated at SunyBiotech (http://www.sunybiotech.com) by using CRISPR–Cas9 methodology. The *linc-133* gain-of-function mutant strain obtained from EMS screening was termed *linc-133(aqz 1)*. This strain contains a mutation in which “A” at 1865 bp of *T09A5.23/linc-133* was replaced with “T”. We used the CRISPR–Cas9 method to recapitulate this mutation *linc-133(syb5664)*, which we termed *linc-133(aqz 3)* in this paper. The following sgRNA primers were used to generate *linc-133(aqz 3)*:

sg1-CCAATAATTTGCCGGCCACCAAC, and sg2-AGACCTGACCAATAATTTGCCGG.

For the deletion allele, 960 bp (1664 bp to 2623 bp) was deleted from *linc-133* to generate *linc-133(syb5619)*, which we termed *linc-133(aqz 2)*, by using CRISPR– Cas9 primers used to generate *linc-133(aqz 2)*:

sg3-GGACTTACTAAATCTTATGACGG, and sg4-CCACATTTCGAACTCTTTCCCTA.

The protein phosphatase of the DAF-18-defective strain *daf-18(D137A)* (PHX6609, *daf-18(syb6609)*) was generated using sg5-CCGAGTCTCGAATTAATGGCTCC. The codon GAT at 459 bp to 461 bp (aspartic acid) of *T07A9.6/daf-18* was replaced with GCG (alanine) using the CRISPR/Cas9 genome editing method.

For the *linc-133* overexpression strains, the gene sequence was amplified from *C. elegans* genomic DNA by using Phusion High-Fidelity PCR (Phusion Master Mix, Thermo Fisher) and cloned into a plasmid (L2528) containing its own promoter.

The primer sequences used for *linc-133* amplification were as follows:

Forward 5’-aattctgcagAATGGGCGGACCCATAGTCA-3’,

Reverse 5’-aattctcgag AACGGTCGAGTTTCAAATAGTGG-3’.

For the *linc-133* gain of function overexpression in AVK cells, the *linc-133* sequence was amplified from the genomic DNA of *linc-133(aqz 3)* strains by using the following primers:

Forward 5’-aattggtaccTTGATCTTAGAGATCGAAAACACGAGAAAGGTG-3’,

Reverse 5’-aattgctagc AGAAATTAAGCCCAATTTATTGAATATTTGGA-3’.

The *flp-1* promoter was amplified from the genomic DNA of N2 worms by using the following primers:

Forward 5’-aattaagctcAATGGGCGGACCCATAGTCA-3’,

Reverse 5’-aattcccggg AACGGTCGAGTTTCAAATAGTGG-3’.

A plasmid with the injection marker *pCFJ90-Pmyo-2::mCherry::unc-54utr* or *pJRK248* was injected into *daf-18(ok480)* worms using standard microinjection methods ^54^. At least three stable lines were generated for each injected strain.

For the plasmids used to construct the *daf-18* rescue strains, please see our previous publication ^2^.

### EMS screening

The *daf-18(ok480)* worms were used to perform genome-wide EMS mutagenesis for forward genetic screening ^15^. In short, more than 800 synchronized L4 stage worms were incubated in a total volume of 4 mL of 50 mM EMS (Sigma) in M9 for 4 h at 20 °C, and approximately 20,000 F2 generation embryos were placed onto NGM plates. The F3 progenies were cultured in M9 to induce L1 arrest and screened for mutants that could survive with normal survival. The selected mutants were backcrossed into *daf-18(ok480)* 3 times. The mutations of interest were identified by using the recently published Sibling Subtraction Method ^24^.

### RNA pulldown

L1-arrested worms were collected and used to prepare total protein extracts. Biotinylated (3’-biotin) *linc-133* probes were designed and synthesized by BGI Tech Solutions Co., Limited (Beijing Liuhe, China). The sequences of the *linc-133* probes are as follows: 5’-TGTGATTTGGAGAAACTGAC-3’; 5’-GTGTGAAAGGACGTCGTCTT-3’; 5’-TTTGAAACACACCCACATCC-3’; 5’-TATGACGGCATTTTCAGCCA-3’; 5’-CGCCACCCTCACCGTTATTA-3’; 5’-CTCTGATCTAGATGCTACCC-3’; 5’-CCGAGTCAGACAAGCTTAAT-3’.

Antisense probes are reverse complements of the above and labeled with biotin. The Pierce Magnetic RNA‒Protein Pull-Down Kit (Thermo Scientific, Cat# 20164) was used to perform an RNA pull down assay to identify the specific proteins interacting with *linc-133*. The procedure was performed according to the manufacturer guidelines. After pull down, equal amounts of protein were separated by electrophoresis on 7.5% polyacrylamide gels (Epizyme Biotechnology, Cat# PG211). Then, the gels were stained using a Fast Silver Stain Kit (Beyotime Biotechnology, Cat# P0017S) according to the manufacturer’s instructions. After silver staining, RNA-binding protein complex-specific bands were cut out and sent to BGI Tech Solutions Co., Limited (Shenzhen, China) for mass spectrometry analysis.

### Western blotting

Protein extraction from L1-arrested larvae was performed by ultrasonic cracking. We measured the protein concentration by using a BCA Protein Assay Kit (Solarbio. Cat#PC0020), separated protein samples via electrophoresis using SDS-containing polyacrylamide gels, and then transferred the separated protein samples onto polyvinylidene fluoride (PVDF) membranes. After blocking with 5% milk in TBST buffer for 1 h, the membranes were incubated at 4 °C overnight with the previously indicated primary antibodies. Afterward, the membrane was washed 3 times with TBST buffer for 10 min each. We incubated the membranes with the corresponding HRP-labeled secondary antibodies for 1 h at room temperature and then washed the membranes 3 times with TBST buffer. Finally, the proteins were visualized using enhanced chemiluminescence (Meilunbio), and the band intensity was determined with a FluorChem M Imaging System (ProteinSimple). The primary antibodies used in the study were anti-ubiquitin (K48 linkage-specific) (Abcam, Cat#ab140601), 14-3-3 polyclonal antibody (Proteintech, Cat#14503-1-AP), and beta tubulin antibody (Abways Technology, Cat#F084106).

### RNA immunoprecipitation (RIP)

RIP was carried out according to previously described methods ^55^. L1-arrested worms were harvested and rinsed twice with ice-cold PBS. The worms were suspended in 1 mL RIP buffer (nuclease-free) containing 50 mM Tris pH 7.4, 150 mM NaCl, 0.5% NP-40, 100 units of RNase inhibitor, 2 mM ribonucleoside vanadyl complex (Sangon Biotech), 1 mM phenylmethylsulfonyl fluoride (PMSF) and 1× protease inhibitor cocktail (MCE). Following brief sonication, worm lysate was centrifuged at 12,000 ×g for 15 min at 4 °C, and the supernatants were precleared with 10 μL of Protein A-coated agarose (Millipore). The precleared supernatants were then divided into two equal parts and incubated with 20 μL 14-3-3 polyclonal antibody and IgG overnight at 4 °C. The Protein A-coated agarose beads were then washed twice with a 3X volume of RIP buffer (nuclease-free) and resuspended in worm lysates containing antibody. The samples were rotated for 1 h and then washed three times with RIP buffer. Then, the beads were spun down at 2000x*g* for 30 s, and the lysate was removed. The crosslinks were reversed by proteinase K buffer (50 mM Tris pH 7.4, 150 mM NaCl, 100 units RNase inhibitor, 0.5% SDS, and 200 μg/mL proteinase K) at room temperature for 30 min before being used for RNA extraction. RNA was subjected to real-time PCR analysis using primers to amplify *linc-133*. The primers used for real-time PCR are listed in Table S2.

### Statistical analyses

All graphed data are presented as the mean ± SEM from at least 3 biological replicates performed in triplicate technical replicates. Statistical calculations were performed using GraphPad Prism 5 (GraphPad Software). The GFP mean intensity was quantified using ImageJ. The mean intensity of Western blotting bands was quantified using the Image J. The relative expression levels of genes were determined by using real-time PCR and the 2^-ΔΔCT^ method. The differences between two groups were analyzed using two-tailed Student’s t test. For multiple comparisons, as in the DAF-16 nuclear localization assay, two-way ANOVA was used to determine the P values. For survival analysis, the mean survival rate was calculated by the Kaplan‒ Meier method. The P values of the difference in overall survival rate were determined using the log-rank test. A P value <0.05 was indicative of statistical significance. Statistical significance values were set as *p < 0.05, **p < 0.01, ***p < 0.001, n.s. = not significant. Further details are provided in the figure legends.

## Supporting information

Supplementary results

## Acknowledgments

We are grateful to Caenorhabditis Genomic Center for providing strains.

## Funding

Henan University (Yellow River Scholar Fund)

Key Scientific Research Project Plan of Henan Province (No. 22A310011)

Henan Province’s key R&D and promotion projects (scientific and technological research) projects (No. 222102310587)

## Author contributions

Conceptualization: SQZ, ZQ

Methodology: SQZ, ZQ

Investigation: WH, FZD, LZ, XY

Visualization: SQZ, WH, FZD, LZ, XY

Funding acquisition: SQZ, ZQ

Project administration: SQZ, ZQ

Supervision: SQZ

Writing – original draft: SQZ

Writing – review & editing: SQZ, ZQ, WH

## Data and materials availability

All data are available in the main text or the supplementary information. The RNA sequencing data can be found in GEO (GSE222091).

