## Supplementary results for "Gain of function in *linc-133* compensates for *daf-18/PTEN* loss and rescues survival"

Fig. S1.

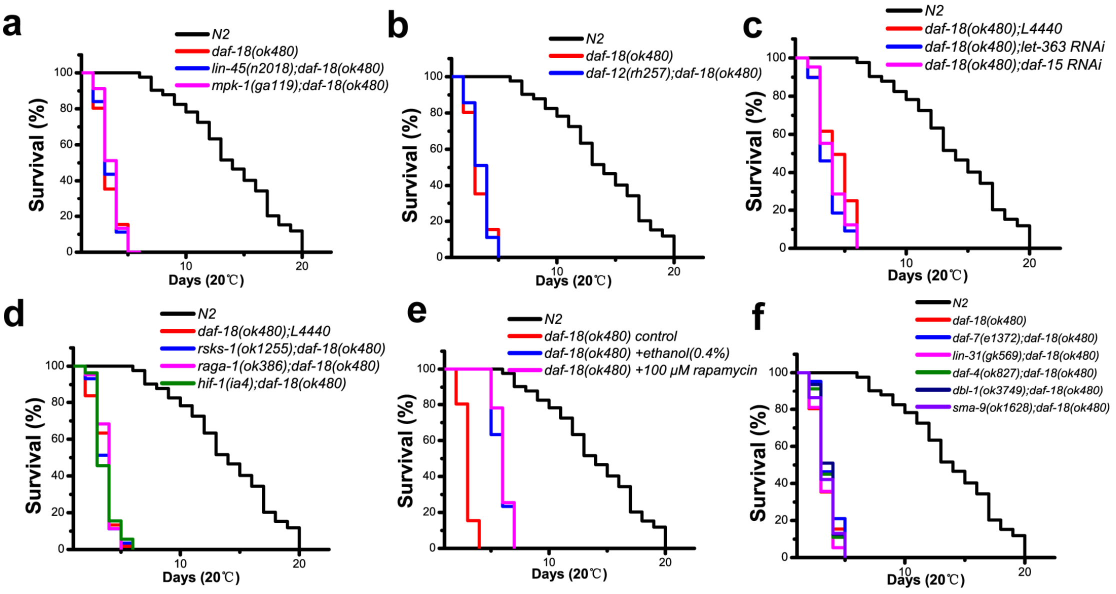


**Figure S1. The effects of mTOR, MAPK, DAF-12 and TGF-beta on survival of *daf-18* L1 arrest worms.**

The survival curves of disrupting MAPK (a), DAF-12 (b), mTOR (c-e) and TGF-beta (f) in L1-arrested worms. Each set of survival experiments was independently repeated at least three times. The mean survival rates were calculated using the Kaplan–Meier method, and P values were determined by using the log-rank test. All the survival data are summarized in Table S1.

**Fig. S2**

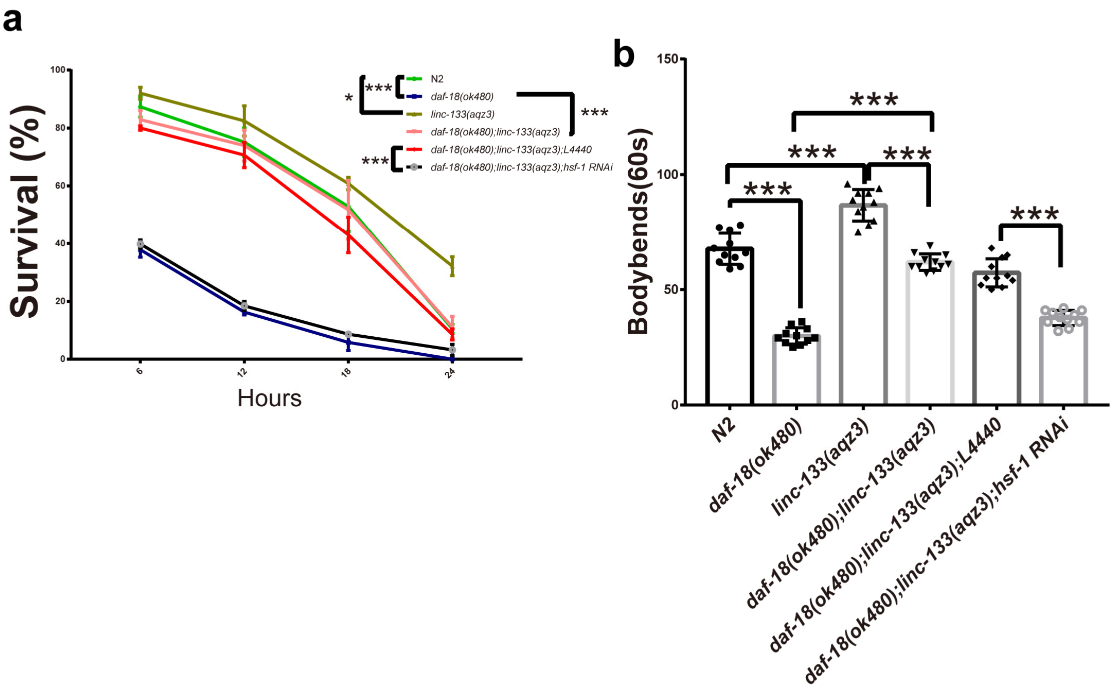


**Figure S2. *linc-133* gain-of-function improves heat stress resistance and mobility.** (a) The effects of *linc-133 aqz3, hsf-1* and *daf-18* on survival of worms under 35℃. (b) the mobility affected by *linc-133 aqz3, hsf-1* and *daf-18* were tested by using thrashing experiments. Animals that were alive but unable to swim scored as paralyzed. Motility assays were carried out by counting body bends over a 60 s interval in M9. Each set of survival experiments was independently repeated at least three times. *:p<0.05, ***: p<0.001.

Table S1. The results of survival during L1 arrest.

| Strains | | Mean life span±SEM (days) | Maximum life span (days) | *P*-value | N |
|---|---|---|---|---|---|
| *daf-18(ok480)* | *L4440* RNAi | 3.91±0.32 | 6 | | >75 |
| | | 4.15±0.14 | 6 | | >59 |
| | | 4.18±0.17 | 6 | | >63 |
| | *let-363* RNAi | 3.95±0.22 | 6 | n.s. | >91 |
| | | 4.20±0.31 | 6 | n.s. | >101 |
| | | 4.13±0.17 | 6 | n.s. | >87 |
| | *daf-15* RNAi | 3.93±0.15 | 6 | n.s. | >65 |
| | | 4.11±0.23 | 6 | n.s. | >60 |
| | | 4.08±0.34 | 6 | n.s. | >68 |
| | *linc-133* RNAi *a* | 4.11±0.06 | 5 | n.s. | ＞90 |
| | | 4.15±0.06 | 5 | n.s. | ＞89 |
| | | 4.09±0.06 | 5 | n.s. | ＞117 |
| | *linc-133* RNAi *b* | 4.08±0.06 | 5 | n.s. | ＞132 |
| | | 4.18±0.06 | 5 | n.s. | ＞140 |
| | | 4.08±0.05 | 5 | n.s. | ＞124 |
| | *linc-133* RNAi *c* | 4.07±0.06 | 5 | n.s. | ＞155 |
| | | 4.11±0.06 | 5 | n.s. | ＞123 |
| | | 4.02±0.05 | 5 | n.s. | ＞108 |
| *daf-18(ok480)* | 4% ethonal | 6.26±0.30 | 7 | | >109 |
| | | 6.15±0.14 | 7 | | >125 |
| | | 6.31±0.09 | 7 | | >136 |
| | 100 μm rapamycin | 6.22±0.17 | 7 | n.s. | >201 |
| | | 6.19±0.25 | 7 | n.s. | >98 |
| | | 6.30±0.24 | 7 | n.s. | >112 |
| *daf-18(ok480)* | control | 4.14±0.06 | 5 | | ＞123 |
| | | 4.21±0.06 | 5 | | ＞87 |
| | | 4.16±0.06 | 5 | | ＞99 |
| | 2 μm AKTi-1/2 | 7.32±0.35 | 10 | ＜0.001 | ＞123 |
| | | 7.15±0.18 | 10 | ＜0.001 | ＞112 |
| | | 7.26±0.32 | 10 | ＜0.001 | ＞98 |
| | *daf-18(ok480)* | 4.14±0.06 | 5 | | ＞101 |
| | | 4.21±0.06 | 5 | | ＞87 |
| | | 4.16±0.06 | 5 | | ＞99 |
| | *daf-18(ok480); mpk-1(ga119)* | 4.01±0.31 | 5 | n.s. | >154 |
| | | 4.15±0.41 | 5 | n.s. | >99 |
| | | 4.12±0.22 | 5 | n.s. | >96 |
| | *daf-18(ok480); rsks-1(ok1255)* | 3.91±0.24 | 6 | n.s. | >69 |
| | | 4.17±0.19 | 6 | n.s. | >66 |
| | | 4.03±0.07 | 6 | n.s. | >73 |

| | | | | | | |
|---|---|---|---|---|---|---|
| | *daf-18(ok480); hif-1(ia4)* | 3.79±0.31 | 6 | n.s. | | >111 |
| | | 4.04±0.15 | 6 | n.s. | | >106 |
| | | 4.06±0.11 | 6 | n.s. | | >95 |
| | *daf-18(ok480); raga-1(ok386)* | 3.89±0.61 | 6 | n.s. | | >96 |
| | | 4.15±0.21 | 6 | n.s. | | >87 |
| | | 4.09±0.31 | 6 | n.s. | | >68 |
| | *daf-18(ok480); daf-4(ok827)* | 4.08±0.14 | 5 | n.s. | | >67 |
| | | 3.99±0.21 | 5 | n.s. | | >85 |
| | | 4.17±0.22 | 5 | n.s. | | >102 |
| | *daf-18(ok480); daf-7(e1372)* | 4.11±0.13 | 5 | n.s. | | >68 |
| | | 4.19±0.09 | 5 | n.s. | | >95 |
| | | 4.08±0.16 | 5 | n.s. | | >96 |
| | *daf-18(ok480); dbl-1(ok3784)* | 4.15±0.12 | 5 | n.s. | | >88 |
| | | 4.08±0.13 | 5 | n.s. | | >80 |
| | | 4.12±0.16 | 5 | n.s. | | >65 |
| | *daf-18(ok480); lin-31(gk569)* | 4.10±0.20 | 5 | n.s. | | >100 |
| | | 4.13±0.13 | 5 | n.s. | | >100 |
| | | 4.19±0.11 | 5 | n.s. | | >108 |
| | *daf-18(ok480); sma-9(ok1628)* | 3.98±0.17 | 5 | n.s. | | >96 |
| | | 4.26±0.50 | 5 | n.s. | | >68 |
| | | 4.18±0.21 | 5 | n.s. | | >60 |
| | *daf-18(ok480); daf-12(rh257)* | 4.11±0.23 | 5 | n.s. | | >64 |
| | | 4.04±0.15 | 5 | n.s. | | >66 |
| | | 4.19±0.28 | 5 | n.s. | | >63 |
| | *daf-18(ok480); mir-235(n4504)* | 3.89±0.07 | 6 | n.s. | | >69 |
| | | 4.15±0.18 | 6 | n.s. | | >79 |
| | | 4.09±0.11 | 6 | n.s. | | >86 |
| | *age-1(m333)* | 18.32±0.05 | 26 | | | >53 |
| | | 17.68±0.81 | 25 | | | >84 |
| | | 18.21±0.63 | 26 | | | >66 |
| | *daf-18(ok480); age-1(m333)* | 7.85±0.09 | 10 | ＜0.001 | | ＞125 |
| | | 7.81±0.09 | 10 | ＜0.001 | | ＞90 |
| | | 7.58±0.10 | 10 | ＜0.001 | | ＞115 |
| | *daf-18(ok480); daf-16(mu86)* | 4.14±0.10 | 5 | vs *daf-16(mu86)* ＜0.001 | | >96 |
| | | 4.25±0.23 | 5 | vs *daf-16(mu86)* ＜0.001 | | >100 |
| | | 4.09±0.17 | 5 | vs *daf-16(mu86)* ＜0.001 | | >125 |
| | *daf-18(ok480); linc-133(aqz1)* | 13.93±0.13 | 19 | ＜0.001 | vs N2＜0.001 | ＞132 |
| | | 13.42±0.13 | 18 | ＜0.001 | vs N2＜0.001 | ＞96 |
| | | 13.79±0.13 | 19 | ＜0.001 | vs N2＜0.001 | ＞121 |
| | *daf-18(ok480); linc-133(aqz2)* | 4.34±0.06 | 5 | n.s. | | ＞115 |
| | | 4.25±0.06 | 5 | n.s. | | ＞136 |
| | | 4.20±0.06 | 5 | n.s. | | ＞101 |
| | | 14.53±0.13 | 18 | ＜0.001 | | ＞123 |

| | | | | | | |
|---|---|---|---|---|---|---|
| | *daf-18(ok480);* | 14.25±0.14 | 17 | ＜0.001 | | ＞100 |
| | *linc-133(aqz3)* | 13.59±0.14 | 17 | ＜0.001 | | ＞110 |
| | *daf-18(ok480);* | 14.68±0.21 | 18 | ＜0.001 | | ＞103 |
| | *daf-16(mu86);* | 14.74±0.36 | 18 | ＜0.001 | | ＞89 |
| | *linc-133(aqz3)* | 13.98±0.85 | 17 | ＜0.001 | | ＞107 |
| *daf-18(ok480)* | Vector | 4.15±0.06 | 5 | | | ＞87 |
| | | 4.28±0.05 | 5 | | | ＞99 |
| | | 4.31±0.06 | 5 | | | ＞119 |
| | *daf-18p::daf-18 D137A(oe)* | 7.35±0.14 | 10 | ＜0.001 | vs N2＜0.001 | >95 |
| | | 7.41±0.28 | 10 | ＜0.001 | vs N2＜0.001 | >90 |
| | | 7.44±0.31 | 10 | ＜0.001 | vs N2＜0.001 | >80 |
| | *daf-18p::daf-18 G174E(oe)* | 10.25±0.42 | 12 | ＜0.001 | vs N2＜0.001 | >66 |
| | | 10.74±0.12 | 12 | ＜0.001 | vs N2＜0.001 | >60 |
| | | 10.39±0.25 | 12 | ＜0.001 | vs N2＜0.001 | >73 |
| | *daf-18p::daf-18 C169S(oe)* | 4.13±0.14 | 5 | n.s. | vs N2＜0.001 | >99 |
| | | 4.21±0.17 | 5 | n.s. | vs N2＜0.001 | >100 |
| | | 4.33±0.22 | 5 | n.s. | vs N2＜0.001 | >125 |
| | *linc-133(oe)* | 4.16±0.06 | 5 | n.s. | | ＞129 |
| | | 4.31±0.05 | 5 | n.s. | | ＞98 |
| | | 4.23±0.05 | 5 | n.s. | | ＞109 |
| | *flp-1p::linc-133 with aqz3(oe)* | 17.35±0.36 | 20 | ＜0.001 | | >153 |
| | | 17.29±0.44 | 20 | ＜0.001 | | >100 |
| | | 17.48±0.27 | 20 | ＜0.001 | | >65 |
| | N2 | 17.97±0.17 | 20 | | | ＞125 |
| | | 17.55±0.16 | 20 | | | ＞114 |
| | | 17.67±0.15 | 20 | | | ＞157 |
| | *daf-16(mu86)* | 7.25±0.12 | 10 | ＜0.001 | | >66 |
| | | 7.23±0.15 | 10 | ＜0.001 | | >60 |
| | | 7.16±0.53 | 10 | ＜0.001 | | >60 |
| | *daf-18(D137A)* | 3.97±0.06 | 5 | ＜0.001 | | ＞118 |
| | | 3.98±0.07 | 5 | ＜0.001 | | ＞97 |
| | | 4.11±0.06 | 5 | ＜0.001 | | ＞108 |
| | *linc-133(aqz1)* | 18.01±0.33 | 20 | n.s. | | ＞105 |
| | | 17.94±0.29 | 20 | n.s. | | ＞110 |
| | | 17.48±0.41 | 20 | n.s. | | ＞95 |
| | *linc-133(aqz2)* | 17.59±0.18 | 20 | n.s. | | ＞78 |
| | | 18.12±0.51 | 21 | n.s. | | ＞74 |
| | | 17.38±0.11 | 20 | n.s. | | ＞95 |
| | *linc-133(aqz3)* | 17.30±0.18 | 20 | n.s. | | ＞97 |
| | | 17.72±0.19 | 20 | n.s. | | ＞82 |
| | | 17.28±0.15 | 20 | n.s. | | ＞69 |
| | *daf-16(mu86);* | 16.35±0.22 | 20 | vs *linc-133(aqz3)* | vs *daf-16(mu86)* | ＞114 |

| | | | | | | |
|---|---|---|---|---|---|---|
| | *linc-133(aqz3)* | | | ＜0.001 | ＜0.001 | |
| | | 16.58±0.16 | 20 | vs *linc-133(aqz3)* ＜0.001 | vs *daf-16(mu86)* ＜0.001 | ＞95 |
| | | 17.01±0.29 | 21 | vs *linc-133(aqz3)* ＜0.001 | vs *daf-16(mu86)* ＜0.001 | ＞143 |
| | *hsf-1(sy441); linc-133(aqz3)* | 6.14±0.22 | 8 | vs *linc-133(aqz3)* ＜0.001 | | ＞78 |
| | | 6.17±0.31 | 8 | vs *linc-133(aqz3)* ＜0.001 | | ＞57 |
| | | 6.12±0.16 | 8 | vs *linc-133(aqz3)* ＜0.001 | | ＞95 |
| | *hsf-1(sy441);* | 7.51±0.41 | 9 | ＜0.001 | | ＞96 |
| | | 7.41±0.19 | 9 | ＜0.001 | | ＞104 |
| | | 7.39±0.18 | 9 | ＜0.001 | | ＞83 |
| N2 | *L4440* RNAi | 16.35±0.11 | 20 | | | ＞95 |
| | | 16.55±0.62 | 20 | | | ＞62 |
| | | 16.17±0.24 | 20 | | | ＞75 |
| | *linc-133* RNAi | 16.05±0.11 | 20 | n.s. | | ＞68 |
| | | 16.87±0.62 | 20 | n.s. | | ＞74 |
| | | 16.43±0.24 | 20 | n.s. | | ＞77 |
| | *par-5 RNAi* | 16.18±0.08 | 20 | n.s. | | ＞85 |
| | | 16.31±0.19 | 20 | n.s. | | ＞69 |
| | | 16.21±0.17 | 20 | n.s. | | ＞72 |
| | *ftt-2 RNAi* | 16.13±0.09 | 20 | n.s. | | ＞69 |
| | | 16.54±0.22 | 20 | n.s. | | ＞102 |
| | | 16.23±0.16 | 20 | n.s. | | ＞79 |
| | *hsp-1 RNAi* | 16.13±0.28 | 20 | n.s. | | ＞69 |
| | | 16.59±0.64 | 20 | n.s. | | ＞94 |
| | | 16.33±0.19 | 20 | n.s. | | ＞70 |
| *daf-18(ok480) ;linc-133(aqz1)* | *L4440* RNAi | 15.89±0.45 | 20 | | | ＞103 |
| | | 15.34±0.32 | 20 | | | ＞97 |
| | | 15.15±0.13 | 20 | | | ＞80 |
| | *linc-133* RNAi | 5.31±0.19 | 7 | ＜0.001 | | ＞76 |
| | | 5.29±0.47 | 7 | ＜0.001 | | ＞84 |
| | | 5.37±0.52 | 7 | ＜0.001 | | ＞93 |
| *linc-133(aqz3)* | *L4440* RNAi | 16.64±0.14 | 20 | | | ＞95 |
| | | 16.37±0.14 | 20 | | | ＞157 |
| | | 16.42±0.14 | 20 | | | ＞171 |
| | *linc-133* RNAi | 15.95±0.29 | 20 | n.s. | | ＞115 |
| | | 16.35±0.77 | 20 | n.s. | | ＞163 |
| | | 16.78±0.43 | 21 | n.s. | | ＞151 |
| *daf-18(ok480) ;linc-133(aqz3)* | *L4440* RNAi | 14.99±0.14 | 18 | | | ＞118 |
| | | 15.01±0.13 | 18 | | | ＞156 |
| | | 14.29±0.13 | 17 | | | ＞74 |

| | | | | | | |
|---|---|---|---|---|---|---|
| | *par-5* RNAi | 9.25±0.41 | 15 | ＜0.001 | | >102 |
| | | 9.11±0.22 | 15 | ＜0.001 | | >100 |
| | | 9.16±0.34 | 15 | ＜0.001 | | >60 |
| | *ftt-2* RNAi | 8.11±0.21 | 14 | ＜0.001 | | >66 |
| | | 8.08±0.23 | 14 | ＜0.001 | | >152 |
| | | 8.34±0.41 | 14 | ＜0.001 | | >114 |
| | *hsf-1* RNAi | 4.24±0.14 | 6 | ＜0.001 | | ＞171 |
| | | 4.35±0.13 | 6 | ＜0.001 | | ＞213 |
| | | 4.58±0.14 | 6 | ＜0.001 | | ＞199 |
| | *hsp-1* RNAi | 11.12±0.13 | 14 | ＜0.001 | | ＞50 |
| | | 11.17±0.13 | 14 | ＜0.001 | | ＞145 |
| | | 10.43±0.11 | 13 | ＜0.001 | | ＞156 |
| | *hsp-70* RNAi | 13.27±0.13 | 16 | n.s. | | ＞198 |
| | | 13.02±0.12 | 16 | 0.0223 | | ＞385 |
| | | 13.41±0.12 | 16 | n.s | | ＞397 |
| | *unc-23* RNAi | 13.77±0.11 | 16 | n.s | | ＞214 |
| | | 13.38±0.11 | 16 | n.s | | ＞217 |
| | | 13.69±0.11 | 16 | n.s | | ＞227 |
| | *hsp-90* RNAi | 13.15±0.11 | 15 | ＜0.001 | | ＞69 |
| | | 13.25±0.09 | 15 | ＜0.001 | | ＞195 |
| | | 13.82±0.10 | 16 | ＜0.001 | | ＞96 |
| | *hsp-110* RNAi | 14.69±0.11 | 17 | n.s. | | ＞105 |
| | | 14.74±0.11 | 17 | n.s. | | ＞93 |
| | | 14.70±0.12 | 17 | n.s. | | ＞143 |
| | *hsp-16* RNAi | 11.98±0.14 | 15 | ＜0.001 | | ＞126 |
| | | 12.14±0.14 | 15 | ＜0.001 | | ＞200 |
| | | 11.97±0.16 | 15 | ＜0.001 | | ＞218 |
| | *cdc-48.1* RNAi | 11.01±0.12 | 13 | ＜0.001 | | ＞274 |
| | | 11.79±0.09 | 14 | ＜0.001 | | ＞245 |
| | | 12.38±0.10 | 15 | ＜0.001 | | ＞197 |
| | *cdc-48.2* RNAi | 11.04±0.10 | 13 | ＜0.001 | vs *cdc-48.1* RNAi ＜0.001 | ＞137 |
| | | 10.42±0.14 | 12 | ＜0.001 | vs *cdc-48.1* RNAi ＜0.001 | ＞92 |
| | | 10.67±0.11 | 12 | ＜0.001 | vs *cdc-48.1* RNAi ＜0.001 | ＞154 |
| *daf-18(D137A) ;linc-133(aqz3)* | *L4440* RNAi | 13.15±0.18 | 19 | | | >90 |
| | | 13.43±0.22 | 19 | | | >93 |
| | | 13.24±0.13 | 19 | | | >60 |
| | *hsf-1* RNAi | 4.86±0.12 | 7 | ＜0.001 | vs *daf-18(D137A)* n.s. | >111 |
| | | 4.92±0.33 | 7 | ＜0.001 | vs *daf-18(D137A)* n.s. | >105 |

| | | | | | | |
|---|---|---|---|---|---|---|
| | | 5.01±0.32 | 8 | ＜0.001 | vs *daf-18(D137A)* n.s. | >162 |
| | *hsp-1* RNAi | 9.05±0.21 | 12 | ＜0.001 | | >75 |
| | | 9.11±0.37 | 12 | ＜0.001 | | >77 |
| | | 9.16±0.19 | 12 | ＜0.001 | | >70 |
| | *cdc-48.1* RNAi | 9.25±0.54 | 14 | ＜0.001 | | >119 |
| | | 9.19±0.33 | 14 | ＜0.001 | | >100 |
| | | 9.33±0.45 | 14 | ＜0.001 | | >124 |
| | *cdc-48.2* RNAi | 8.53±0.19 | 12 | ＜0.001 | vs *cdc-48.1* RNAi ＜0.001 | >96 |
| | | 8.41±0.23 | 12 | ＜0.001 | vs *cdc-48.1* RNAi ＜0.001 | >80 |
| | | 8.22±0.37 | 12 | ＜0.001 | vs *cdc-48.1* RNAi ＜0.001 | >105 |

Mean lifespans were calculated by using Kaplan-Meier method; *P*-values compared with Controls were determined using the log-rank test; n.s., no significant difference; SEM, standard error of the mean; N, number of worms counted. L4440: RNAi control. Vector: empty expression vector as a control.

Table S2. Primers and RNAi used in this work.

| **Methods** | **Gene name** | **Primer sequence** |
|---|---|---|
| Worm RNAi | *linc-133 a* | F:5’-AATT AGATCT ACGAGAAAGGTGTACGTCCTG-3’ |
| | | R:5’-AATT GCTAGC ATATTTGGCTTGCGGGGGCT-3’ |
| | *linc-133 b* | F:5’-AATT AGATCT AGAGACAGCAAGAAGAAGTGAGT-3’ |
| | | R:5’-AATT GCTAGC GGAGAGAGGCATCGGATGAA-3’ |
| | *linc-133 c* | F:5’-AATT CCCGGG CGATTCTCCCTGATCATCGT-3’ |
| | | R:5’-AATT AAGCTT TGATCAAAATGATGCGGAAA-3’ |
| | *let-363* | F:5’- AATT TCTAGA AGGCATAGACCACAAGCAGT-3’ |
| | | R:5’- AATT AAGCTT CCAACGAGTTCAAAGTGGCA -3’ |
| | *daf-15* | F:5’-AATT AGATCT AGTGATTGGGCGATTCATGC-3’ |
| | | R:5’-AATT CTCGAG CTCCTGGTCCTGTCTGTTGT -3’ |
| | *hsp-16.11* | Horizon RNAi library (Horizon Discovery Ltd) |
| | *hsp-16.48* | Horizon RNAi library (Horizon Discovery Ltd) |
| | *par-5* | Horizon RNAi library (Horizon Discovery Ltd) |
| | *ftt-2* | Horizon RNAi library (Horizon Discovery Ltd) |
| | *hsp-1* | Horizon RNAi library (Horizon Discovery Ltd) |
| | *hsp-70* | Horizon RNAi library (Horizon Discovery Ltd) |
| | *cdc-48.1* | Horizon RNAi library (Horizon Discovery Ltd) |
| | *cdc-48.2* | Horizon RNAi library (Horizon Discovery Ltd) |
| | *unc-23* | Horizon RNAi library (Horizon Discovery Ltd) |
| | *hsp-90* | Horizon RNAi library (Horizon Discovery Ltd) |
| | *hsp-110* | Horizon RNAi library (Horizon Discovery Ltd) |
| | *hsf-1* | Horizon RNAi library (Horizon Discovery Ltd) |
| | *linc-133 1* | 5’-TTTCTCGTGTTTTCGATCTC-3’ |
| | *linc-133 2* | 5’-GCAATCTTTGTCTTCAGGAC-3’ |
| | *linc-133 3* | 5’-GAGTTGTCAATCATCGGGAG-3’ |
| | *linc-133 4* | 5’-GTCAGTTTCTCCAAATCACA-3’ |
| | *linc-133 5* | 5’-TCTACTTTGAATGCCCATTT-3’ |
| | *linc-133 6* | 5’-GATACGCAGACTGGTCATTC-3’ |
| | *linc-133 7* | 5’-ATGTTTCTGGTTCAGTTGTC-3’ |
| | *linc-133 8* | 5’-AAGACGACGTCCTTTCACAC-3’ |
| | *linc-133 9* | 5’-AATTTAGAAGTGACCCTTCC-3’ |
| | *linc-133 10* | 5’-CACATTCATCCACTTTGCAA-3’ |
| | *linc-133 11* | 5’-GGTGACCAATGCTCAATTTC-3’ |
| | *linc-133 12* | 5’-ATTTGCCAGAATCATTGCGA-3’ |
| | *linc-133 13* | 5’-GTAGCGGTTTTACGTTTTGA-3’ |
| | *linc-133 14* | 5’-AGCCGATAGGGAGGAGTAAA-3’ |
| | *linc-133 15* | 5’-CTCACTTTCTGTTCCATATT-3’ |
| | *linc-133 16* | 5’-CTTGCTGTCTCTTTCTGAAT-3’ |

| | | |
|---|---|---|
| smFISH *linc-133* probe | *linc-133 17* | 5'-GGATGTGGGTGTGTTTCAAA-3' |
| | *linc-133 18* | 5'-ACACACATTGCCCTACTAAA-3' |
| | *linc-133 19* | 5'-GTGGTTTATCTCATGCATGA-3' |
| | *linc-133 20* | 5'-CCGATCGGATAACTAGTTCA-3' |
| | *linc-133 21* | 5'-CTTCCATTTTGGCAGTAATT-3' |
| | *linc-133 22* | 5'-TGGCTGAAAATGCCGTCATA-3' |
| | *linc-133 23* | 5'-TCAGGTCTTCTGTGTAGATT-3' |
| | *linc-133 24* | 5'-TTGCGAAGAACCTTTCCTAC-3' |
| | *linc-133 25* | 5'-AGGGGCACACGCATAATATA-3' |
| | *linc-133 26* | 5'-GTATGAGGTTGAGGGAGAGGG-3' |
| | *linc-133 27* | 5'-ATGTACCGCCTGGAGAAATA-3' |
| | *linc-133 28* | 5'-AAGGAGAGAGGCATCGGATG-3' |
| | *linc-133 29* | 5'-TAATAACGGTGAGGGTGGCG-3' |
| | *linc-133 30* | 5'-CGGGCATGTTCATACAACTT-3' |
| | *linc-133 31* | 5'-AGTTTCCGTGAAACGGATGG-3' |
| | *linc-133 32* | 5'-GTAGTTCACACTTTCACTTC-3' |
| | *linc-133 33* | 5'-TTCACCAATATCTTTCGTGC-3' |
| | *linc-133 34* | 5'-TCCGTGTAATTGTGATGACC-3' |
| | *linc-133 35* | 5'-CTCGTTTTTCAGGTTTCATT-3' |
| | *linc-133 36* | 5'-AGGCAGTAAGCCAATTGTTT-3' |
| | *linc-133 37* | 5'-AATGAGAGGGACGCGGAACG-3' |
| | *linc-133 38* | 5'-GAGACGAAAGTAGGCGGAGT-3' |
| | *linc-133 39* | 5'-CGGGCATGAATGTGTTTGTA-3' |
| | *linc-133 40* | 5'-TAAGCGATGATGGTGGTGG-3' |
| | *linc-133 41* | 5'-ATTCAGGCGATAGGCATATA-3' |
| | *linc-133 42* | 5'-GGGTAGCATCTAGATCAGAG-3' |
| | *linc-133 43* | 5'-ACCGACGAGCAACATGATTT-3' |
| | *linc-133 44* | 5'-AAAAGGATGGAGTGTGCTCC-3' |
| | *linc-133 45* | 5'-GTTCGGTTACAAGAGCTCAT-3' |
| | *linc-133 46* | 5'-GGAGCTCATGATTTGTTCTT-3' |
| | *linc-133 47* | 5'-ATTAAGCTTGTCTGACTCGG-3' |
| | *linc-133 48* | 5'-ACTCAGAGCTTGTTTTCTGA-3' |
| Real-time | *cdc-42* | F:5'-CTGCTGGACAGGAAGATTACG-3' |
| | | R:5'-CTCGGACATTCTCGAATGAAG-3' |
| | *linc-133* | F:5'-CGATTCTCCCTGATCATCGT-3', |
| | | R:5'-CGTGAAACGGATGGATTACC-3' |
| | *hsp-16.11* | F:5'-CTCGCAGTTCAAGCCAGAAG-3' |
| | | R:5'-CCAACATCAACATCTTCGGGT-3' |
| | *hsp-16.48* | F:5'-CTCATGCTCCGTTCTCCATT-3' |
| | | R:5'-GAGTTGTGATCAGCATTTCTCCA-3' |
| | *hsp-70* | F:5'-AGCCCGTTGTTGAGGTTGAA-3' |
| | | R:5'-CCCGTACAGAATGCCCAAGT-3' |
| | *hsp-90* | F:5'-TTCCCGCATGAAGGAGAACC-3' |

| | | |
|---|---|---|
| PCR | | R:5'-CCTTCCTTGGTGACGGAGAC-3' |
| | *hsp-110* | F:5'-GCAATTGCTCTAGCCTACGG-3' |
| | | R:5'-AAGCAACCAATGAAGCCTGT-3' |
| | *unc-23* | F:5'-CTGAGGTACCCGAGAGACCA-3' |
| | | R:5'-GGTTCTCATCACCCTTTCCA-3' |
| | *hsp-1* | F:5'-CTGCGTGGGAGTTTTCATGC-3' |
| | | R:5'-ACTGAACTGCTGGATCGTCG-3' |
| | *let-363* | F:5'-TGGTTCGAGAGGCAAGTCTT-3' |
| | | R:5'-TCGCCTGCACATTGTTATCG-3' |
| | *daf-15* | F:5'-GTCTAACCACGCCTATCCGA-3' |
| | | R:5'-GCACAGTTCTTCGTTCAGCA-3' |
| | *par-5* | F:5'-GCTAAGGACAAGATGCAGCC-3' |
| | | R:5'-CGTGCTCTGGAGTGTTCAAG-3' |
| | *ftt-2* | F:5'-ACCCTCATCATGCAGCTTCT-3' |
| | | R:5'-TCTGTCTCGTTGGCATCAGT-3' |
| | *cdc-48.1* | F:5'-CCAAAACGCGAGAAGACCAA-3' |
| | | R:5'-TGGACGATTTGTTGCAGCAA-3' |
| | *cdc-48.2* | F:5'-AAAGGAGCGTCAGGACAGAA-3' |
| | | R:5'-CGAACTTCATTGCCTCCTCG-3' |
| | *hsf-1* | F:5'-ATGACTCCACTGTCCCAAGG-3' |
| | | R:5'-TGCGTCTGTTCGTTAACCTG-3' |
| | *sod-3* | F:5'-TCGGTTCCCTGGATAACTTG-3' |
| | | R:5'-CATAGTCTGGGCGGACATTT-3' |
